# The RNA binding ubiquitination ligase MEX3B regulates bFGF-dependent neuronal proliferation

**DOI:** 10.1101/2024.06.28.601280

**Authors:** Kamakshi Garg, Gourav Sharma, Sarbani Samaddar, Sourav Banerjee

## Abstract

**Background:** E3 ubiquitin ligases, integral components of the proteasomal degradation cascade, are critical for regulating the cellular proteome via canonical proteasome-mediated protein degradation; however, the non-canonical functions of these ligases in neuronal development are poorly understood. Our study focuses on a special class of E3 ubiquitin ligases known as RNA Binding Ubiquitin Ligases (RBUL) that harbour RNA-binding domains; allowing them to acquire all the properties of RNA-binding proteins (RBPs) and regulate transcriptional or post-transcriptional changes associated with the control of gene expression in cellular phenotypes. We aim to identify one such RUBL in the context of the highly dynamic yet stringently controlled process of neural proliferation and neural fate determination.

**Results:** MEX3B protein is a member of the MEX3 family and a part of the RBUL class of E3 ligases. It is differentially expressed in Neural Progenitor Cells (NPCs) upon differentiation. Mex3b RNA and protein were found to have temporally opposing expression patterns in presence of basic fibroblast growth factor (bFGF), a key signalling protein involved in neuronal proliferation. MEX3B is required for maintenance of the proliferative state of NPCs, whereas, its knockdown promotes transition from proliferative to differentiation state even in presence of bFGF that restricts differentiation. Furthermore, the knockdown of MEX3B protein results in the appearance of morphological hallmarks associated with early stages of neuronal differentiation including increase in neurite length and complexity. MEX3B interacts with the pro-proliferative transcription activator REST and the long non-coding RNA, *HOTAIR*. The study suggests the existence of a bFGF-dependent, combinatorial axis involving Mex3b, REST and *HOTAIR*, for the maintenance of NPC proliferative states.

**Conclusion:** MEX3B, containing RNA binding motifs, is a unique E3 ligase that is necessary for bFGF-dependent proliferation. Mex3b protein invokes its non-canonical function of an RNA binding protein to form a tripartite complex with the transcription activator REST and *HOTAIR* lncRNA to define the proliferative state of NPCs. The study highlights a unique feature of special E3 ligases in neuronal proliferation during brain development that was previously overlooked.

## Background

Amongst the numerous post-translational modifications regulating the cellular proteome; ubiquitination stands out because of its ability to control protein turnover rates by marking proteins for degradation. A tripartite cascade of enzymes is involved in the ubiquitination process; the ubiquitin activating enzyme E1, the ubiquitin-conjugating enzyme E2, and the E3 ubiquitin ligase. E3 ligases constitute the last step of the ubiquitination process, wherein they catalyse the transfer of ubiquitin to specific substrates The vast number (∼1000) and diversity of E3 ligases [1–4] present in eukaryotes allow them incredible versatility in the regulation of protein fates; including protein stability [4–6], the subcellular distribution of proteins [7] and protein degradation either *via* the proteasomal machinery [8, 9] or by autophagy [10–12]. Such control over the cellular proteome bestows E3 ligases with the ability to influence key processes such as cell cycle regulation [13–15], cell migration [16, 17], apoptosis [18]; and overall mammalian development [19].

Within neurons, E3 ligases influence the neuronal proteome and they are known to control various aspects of neuronal development; including neurogenesis [20–22], neurite development [23–25]; regulation of neuronal morphogenesis [26–28], synapse formation [29–32]; synapse pruning [33, 34] and in both Hebbian [35] and homeostatic [36, 37] [31] forms of synaptic plasticity; among others. Aberrations in E3 ligase functions have been linked to disorders involved in neuronal development [38–40] as well as neurodegeneration [41–44].

Subsets of E3 ligases have alternate identities. This is attributed to the diverse functions of different protein domains present in addition to their standard RING, HECT, or RBR (Ring between Ring) domains responsible for their function as ubiquitin ligases [3, 45]. The “moonlighting” function of these E3 ligases provides an additional level of control over gene expression within the cellular compartments; this is especially important for E3 ligases which contain RNA binding domains (RNA binding Ubiquitin Ligases or RUBLs) [45–48]; for e.g. the TRIM family of E3 ligases [46, 48, 49], and Roquins [47]. Within neurons, the RNA-binding ability of these E3 ligases provides them with additional control of the cellular proteome by dynamically regulating transcription, post-transcriptional and translation paradigms [21–25]. Specially during neurogenesis, the temporal expression of genes is very tightly regulated, and there is heightened necessity for the participation of RBULs, as they are capable of coupling RNA metabolism and protein turnover [45, 49]. This, in combination with reports of them being involved in various stages of neuronal development [21–29], led us to investigate how E3 ligases regulate the transition of neuronal progenitor cells from a state of proliferation to that of differentiation. Specifically, we wanted to understand whether the RNA-binding abilities of development-associated E3 ligases provide them with mechanistic advantages to control the expression of genes relevant to driving the proliferation-differentiation balance at the onset of neuronal development.

In this study, we identify that a RNA binding E3 ligase, MEX3B, promotes the maintenance of proliferative states in NPCs. Mex3b belongs to the Mex3 family of RBULs [50, 51], that show different levels of expression when NPCs undergo transition from proliferative to differentiated states. No studies so far document the involvement of Mex3b in determining the fate of cortical progenitor cells. We find that Mex3b aids in maintaining sustained transcription and post-transcriptional modification of pro-proliferative genes. Despite the influence of a pro-proliferative cue like bFGF, loss of function of MEX3B protein results in the partial gain of neuronal identity in progenitor cells, including the appearance of neurites, increase in neurite lengths and the enhanced expression of genes that aid neuronal differentiation. In the presence of bFGF, MEX3B associates with the *REST* mRNA and the long noncoding RNA (lncRNA) *HOTAIR via* its RNA-binding domains to create a coordinated RBP-RNA complex; indicating that the tripartite MEX3B-HOTAIR-*REST* complex may play a role in the maintenance of the proliferative states of NPCs. Taken together, this study showcases how the non-canonical RNA binding properties of neuronal E3 ligases (RBULs) allow them to determine the fate of neuronal progenitors during development.

## Results

### Identification of differentially expressed E3 ligases in neural progenitor cells during the transition from proliferative to differentiated states

To identify an E3 ubiquitin ligase which is expressed differentially in neural progenitor cells (NPCs) tending to differentiation we cultured NPCs with bFGF (basic fibroblastic growth factor) and without bFGF from E11.5 CD1 embryos (Fig.1A). The expression of proliferation and differentiation markers was checked in both plus and minus bFGF conditions at DIV 5 to validate the potency of our system (Fig.1B-C). The relative expression of the RNA of all the proliferation markers viz. Ki67 (0.5422 ± 0.1530 fold decrease, p=0.0165), Nestin (0.2298 ± 0.0610 fold decrease, p=0.0328), REST (0.4462 ± 0.07872 fold decrease, p=0.0109), Notch was found to be reduced (0.5354 ± 0.08107 fold decrease, p=0.0006) in NPCs grown in the presence of bFGF vs. those grown in its absence (Fig. 1B). Neurobasal media containing B27 supplements and antibiotics was used as the culturing medium throughout [52, 53]. Similarly, the relative expression of the RNA of all the differentiation markers such as Homer (4.872 ± 1.330 fold increase, p=0.0105), GFAP (17.09 ± 5.847 fold increase, p=0.0329), β-tubulin III (5.567 ± 1.830 fold increase, p=0.022 8), Snap25 (11.25 ± 3.178 fold increase, p=0.0122), Shank (3.759 ± 1.141 fold increase, p=0.0216), NeuroD1 (9.875 ± 2.627 fold increase, p=0.0094), NeuN (7.768 ± 2.925 fold increase, p=0.0451), Nav1.2F (9.690 ± 3.315 fold increase, p=0.0265) were elevated in NPCs grown without bFGF (Fig. 1C). Our results indicated that the development of NPCs under the influence of bFGF is a valid reductionist model for characterizing E3 ligases that regulate neuronal development, and that the withdrawal of bFGF from the culture media drives differentiation. Next, we performed microarrays from total RNA extracted from NPCs grown with (plus-bFGF), or without bFGF (minus-bFGF) to identify differentially expressed E3 ubiquitin ligase genes, followed by PANTHER GO analysis on all the E3 ubiquitin ligases. Numerous E3 ligases were found to be differentially expressed in the microarray, (Fig.1D and S1A), amongst which, members of the Mex3 family and the Ube3 family showed maximum differential expression. GO terms that were significantly enriched in the GO ‘Molecular function’ class included the GO term ‘binding’ (Fold enrichment= 1.28; FDR=0.0498) (Fig.1F). Further investigation of the PANTHER protein classes found in the GO term ‘binding” led us to discover that all members of the Mex3 family of proteins were common to “binding” and a special protein class termed “RNA metabolism” (Fig. 1G). We also isolated the list of RNA binding proteins reported so far [54] and compared with the entire set of E3 ligases in the microarray to find a subset of E3 ligases with RNA binding properties. 17 E3 ligases were identified as RNA binding proteins as well; hence, they are a part of the RBUL (RNA Binding Ubiquitin class of E3 ligases) family (Fig. 1E and Table. S1). All MEX3 family members were present among them. Thus, the MEX3 family of proteins was our topmost contender for delineating how E3 ligases with RNA binding domains influence neuronal development. We selected members of the MEX3 family for further validation of the microarray data using qPCR. βIII-tubulin was used as a differentiation marker (Fig.2A). A significant upregulation of βIII-tubulin (16.65±4.773 fold increase, p<0.0001) and *Mex3b* (9.107±0.0129 fold increase, p=0.0129) was observed while *Mex3a* (0.2029±0.1203 fold decrease, p=0.9532), and *Mex3d* (2.167±0.8618 fold increase, p=0.5326) did not show any significant change in NPCs grown in plus-bFGF versus in NPCs grown in minus-bFGF conditions. The results identified that transcript levels of *Mex3b*, which belongs to a special family of E3 ubiquitin ligases with RNA-binding domains, are higher in differentiating NPCs. Hence, we selected Mex3b to further study and characterize in neural development paradigms.

**Figure 1:**
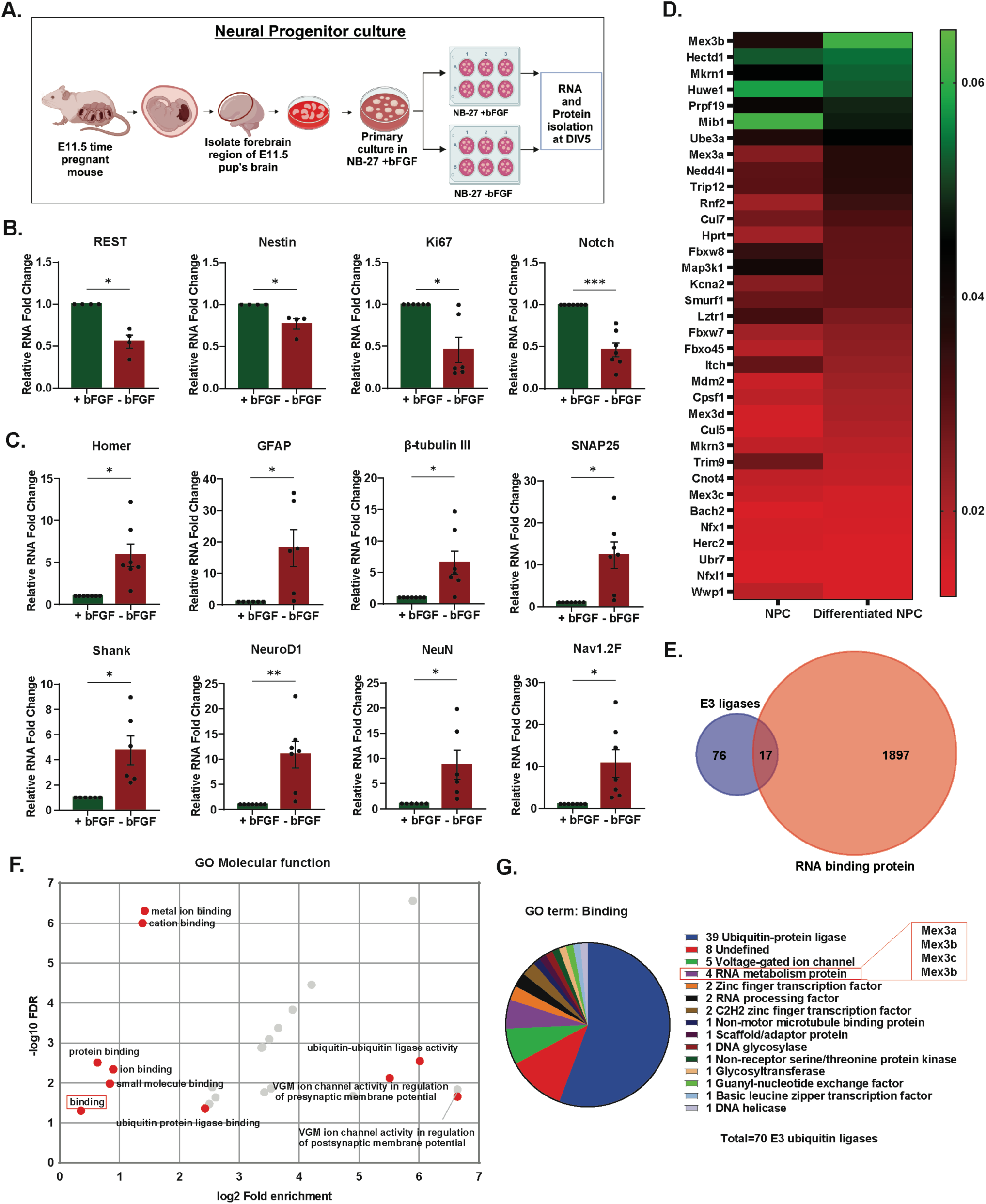
RNA binding E3 Ubiquitin ligase Mex3b is differentially expressed in NPC. **A.** Schematics for Neural progenitor culture protocol. **B.** Differential expression of proliferation markers in differentiating vs proliferating NPCs obtained by qRT-PCR from n=4-7, *p<0.05, ***p<0.001 by Unpaired t test with Welch’s correction. **C.** Differential expression of differentiation markers in differentiating vs proliferating NPCs obtained by qRT-PCR from n=6-7, *p<0.05, **p<0.01 by Unpaired t test with Welch’s correction. **D.** Heatmap representation of expression of a subset of Microarray for E3 Ubiquitin ligases in proliferating and differentiating NPCs from n=2. Organized in order of decreasing expression in NPCs. **E.** Venn diagram showing the overlap between E3 ubiquitin ligases used in microarray and the RNA binding proteins reported in previous study **F.** GO analysis for molecular function from all the E3 ubiquitin ligases used in microarray data (Fig.S1-A.). **G.** Pie chart representation of all the Panther protein class under GO term: ‘Binding’ highlighting RNA metabolism protein. All qRT-PCR data is shown as Mean±SEM.

**Figure 2:**
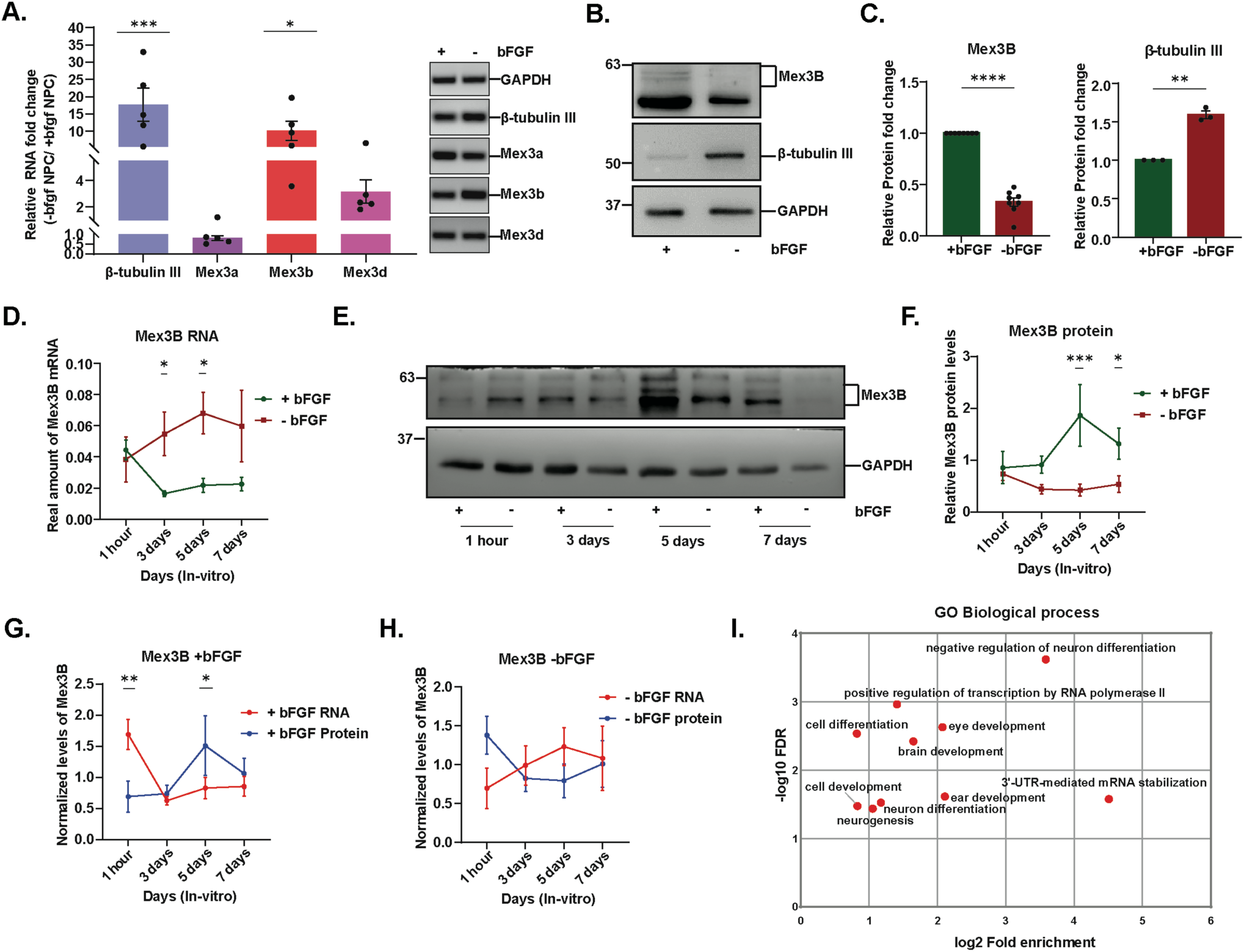
Inverse expression of Mex3b RNA and Protein. **A.** qPCR validation of MEX3 family of E3 ubiquitin ligases represented in the microarray data in NPCs from n=5, *p<0.05, ****p<0.0001 by One-way ANOVA and Fisher’s LSD and Representation of qPCR data from histogram on 2% agarose gel. **B-C.** Western blot quantification of MEX3B and β-tubulin III protein levels in differentiating vs proliferating NPCs from n=8 and 3, ****p<0.0001, **p<0.01 respectively by Unpaired t test with Welch’s correction. **D.** qRT-PCR based comparison of Mex3B RNA at different time point in NPCs in plus and minus-bFGF condition, n=6, *p<0.05 by Two-way ANOVA with Fisher’s LSD. **E-F.** Western blot quantification of MEX3B protein at different time point in NPCs in plus and minus-bFGF condition, n=6, *p<0.05, ***p<0.001 by Two-way ANOVA with Fisher’s LSD. **G-H.** Comparative analysis of Mex3b protein and RNA in NPC in plus-bFGF across the time points, n=6, *p<0.05, **p<0.01 by Two-way ANOVA with Fisher’s LSD. **H.** Comparative analysis of Mex3b protein and RNA in NPC in minus-bFGF across the time points, n=6, all not significant by Two-way ANOVA with Fisher’s LSD. All the data is shown as Mean±SEM. **I.** GO analysis for Biological process from Mex3B targets obtained from Microarray data from previous studies.

### Opposite expression patterns of Mex3b mRNA and protein observed during neuronal proliferation

To test the functional properties of *Mex3b* gene in NPCs, it is imperative to test the effect of NPC differentiation on MEX3B protein levels. Hence, MEX3B protein expression was tested in plus-bFGF and minus-bFGF conditions. To our surprise, MEX3B protein was observed to be downregulated (0.6714 ± 0.04210 fold decrease, p=<0.0001) in NPC grown in NB-27 minus-bFGF versus the NPC grown in NB-27 plus-bFGF which is in complete contrast to RNA expression. βIII-tubulin (0.5894 ± 0.04963 fold increase, p=0.007) was used as a differentiation marker for NPCs that showed upregulation in the differentiation state and GAPDH was used for internal normalization (Fig. 2B-C). To justify this discrepancy in the RNA and protein expression we came across a study [55] that talks about the posttranscriptional autoregulation of *Mex3b* mRNA by binding of its protein to its 3’UTR region. To test the possibility of this autoregulation in our system we further tested the levels of Mex3b protein and RNA at different time points (1hr, 3 days, 5 days and 7 days) in NPC culture and checked the comparative change in their expression patterns. We observed that, in –bFGF conditions, *Mex3b* RNA levels increase from 1hr (0.0385±0.014431) to DIV3 (0.05481±0.014009) and further till DIV5 (0.06813±0.01332) but start decreasing from DIV5 to DIV7 (0.05983±0.022922); while in plus-bFGF condition its levels decrease from 1hr (0.04447±0.006388) till DIV3 (0.01647±0.001759) and then become stable at DIV5 (0.02177±0.004396) and DIV7 (0.02252±0.004232). *Mex3b* mRNA levels were significantly higher in minus-bFGF condition than in plus-bFGF condition at DIV3 (0.03835±0.01767 fold increase, p = 0.0464) and DIV5 (0.04636±0.01767 fold increase p=0.0192) (Fig. 2D). Mex3B protein levels in NPCs grown in plus-bFGF condition remain stable from 1hr (0.855895±0.310668) till DIV3 (0.854601±0.164757) and then peaks at DIV5 (2.662656±0.592516). A little decrease was observed at DIV7 (1.397468±0.298238); while in minus-bFGF condition MEX3B protein levels remain stable throughout. Furthermore, MEX3B levels were significantly higher in +bFGF condition as compared to –bFGF condition at DIV5 (1.444±0.3310 fold increase, p= 0.0006) and DIV7 (0.7784±0.0328 higher p=0.0328) (Fig.S2E-F). Since we observed both protein and RNA changing at only DIV 5 we focused all our future studies at this time point. We further noticed that the presence of bFGF has significant effect on the upregulation of protein levels at DIV5 from DIV3 (0.9535±SEM higher, p<0.05) (Fig. 2E and F).

Additionally, we compared the normalized RNA and protein levels in both plus and -bFGF conditions. In the presence of bFGF, *Mex3b* RNA levels (1.690511±0.242811) are significantly higher (0.99963±0.2692 fold increase, p=0.0021) compared to MEX3B protein levels (0.690881±0.250772) at 1hour time point, but become equal at DIV3 (Protein: 0.73766±0.132993 fold change, RNA: 0.626001±0.066851 fold change). Further on, at DIV5, MEX3B protein levels increase drastically (1.507355±0.478281 fold change) but mRNA levels (0.827505±0.167117 fold change) remain constant; resulting in the significant increase in protein levels as compared to RNA levels (0.67985±0.2692 higher, p=0.0233). At DIV 7, MEX3B protein and *Mex3b* RNA levels again appear to attain equilibrium (Protein: 1.064104±0.240739 fold increase; RNA: 0.855982±0.160872 fold increase). Overall, in the presence of bFGF, protein and RNA levels of Mex3b display inversely correlated patterns of expression. In the absence of bFGF, there were no significant changes between Mex3b RNA and protein levels (Fig.2G-H). This implies the role of bFGF in Mex3b regulation, corroborating to previous observations[55].

To check the physiological relevance of MEX3B protein and *Mex3b* RNA having opposite expression patterns, we assessed both protein and RNA levels in the cortical tissue of mouse at different developmental time points. In the developing mouse brain there are temporally regulated crosstalks between various proliferative and pro-differentiation cues and Mex3b expression patterns could be influenced by many factors other than bFGF. We found that in complete contrast to our findings in NPCs, *Mex3b* RNA levels were significantly higher at E11.5 (1.794±0.3824 higher; p=<0.0001), E15 (1.137±0.3824 higher; p=0.0043), P0 (0.8597±0.3824; p=0.0285) and then become significantly lower at P7 (1.428±0.3824 lower; p=0.0004) and P14 (1.91±0.3824; p=<0.0001) with respect to MEX3B protein. mRNA and protein levels of Mex3b seem to equilibrate at P21 and P28 (Fig. SA-D). This lends further credence to our hypothesis that elevated MEX3B levels in proliferating NPCs were a result of bFGF’s influence. Since the mRNA and protein expression changes inversely to each other both in culture and *in vivo*, it becomes crucial to understand, how MEX3B protein is functioning in NPCs? What is the role of MEX3B in proliferating NPCs? To get an insight into the plausible role of MEX3B we did a Gene ontology analysis on Mex3b targets obtained from a previous study from Masanori Taira’s Lab [55]. GO analysis using PANTHER shows that Mex3b is affecting several important biological processes like Neurogenesis, brain development, eye development, ear development, 3’UTR-mediated mRNA stabilization and positive regulation of transcription by RNA polymerase II, etc (Fig. 2I). Prompted by these observations we further investigated the role of Mex3b in NPCs.

### MEX3B knockdown specifically reduces the transcriptional expression of neural proliferation markers in neuronal progenitors

We wanted to investigate whether MEX3B is capable of controlling proliferation or differentiation stages in NPCs. To this end, two shRNA constructs were cloned against *Mex3b* into the pLVTHM plasmid, a Lentiviral vector. Knockdown efficacy of the two constructs was assessed in the NPC culture system and successful knockdown was evaluated for *Mex3b* RNA by qRT-PCR and MEX3B protein by Western blot. Percentage decrease of *Mex3b* RNA for MEX3B RNAi (shRNA construct#1(plus-bFGF 0.4503±0.08644 fold decrease; p=0.0002);(minus-bFGF 0.5448±0.1351 fold decrease; p=0.0012) and shRNA construct#2 (plus-bFGF 0.5488±0.1059 fold decrease; p=0.0002);(minus-bFGF 0.4792±0.1471 fold decrease; p=0.0062)) was observed relative to Control RNAi. Percentage decrease of MEX3B protein for in RNAi samples (Mex3b shRNA construct#1 (plus-bFGF 0.5654±0.09257 fold decrease; p=0.0003); (minus-bFGF 0.544±0.164 fold decrease; p=0.009) and for Mex3b shRNA construct#2 (plus-bFGF 0.6917±0.09999 fold decrease; p=0.0001); (minus-bFGF 0.5793±0.164 fold decrease; p=0.0064)) was observed relative to Control RNAi (Fig.3A-C). Once Mex3b knockdown was confirmed both at transcript and protein level, the lentivirus containing a plasmid expressing shRNA against *Mex3b* and EF1α-GFP was added into the NPCs. At DIV5 the infected NPCs transcriptome was evaluated for differential expression of neural proliferation and differentiation markers in both with and without bFGF conditions. Decreased expression of Ki67 (plus-bFGF: Mex3b shRNA construct #1: 0.5648±0.1514 fold decrease; p=0.0058; construct #2: 0.6758±0.1635 fold decrease; p=0.0033);(minus-bFGF: Mex3b shRNA construct#1: 0.6387±0.09614; p=<0.0001; construct#2: 0.5305±0.1177; p=0.0007)), REST((plus-bFGF: Mex3b shRNA construct #1: 0.3282±0.08392; p=0.0045; construct #2: 0.5222±0.09065; p=0.0004); (minus-bFGF: Mex3b shRNA construct #1: 0.6549±0.1791; p= 0.0106; construct #2: 0.6002±0.1791; p=0.0154)) and Nestin ((plus-bFGF: Mex3b shRNA construct #1: 0.317±0.1041; p=0.0159; construct #2: 0.3351±0.1124; p=0.0176); (minus-bFGF: Mex3b shRNA construct #1: 0.5017±0.07193; p=0.0004; construct #2: 0.4187±0.07193; p=0.0011) in Mex3b shRNA construct #1 and #2 with respect to control shRNA was observed in both conditions (Fig.3D-F).Expression of differentiation markers did not show any significant change at the transcript level, both with and without bFGF conditions (Fig.S3B-C). Mex3b seems to have transcriptional or post-transcriptional control on the transcripts of proliferation markers but not on differentiation markers in NPCs in the presence of bFGF.

**Figure 3:**
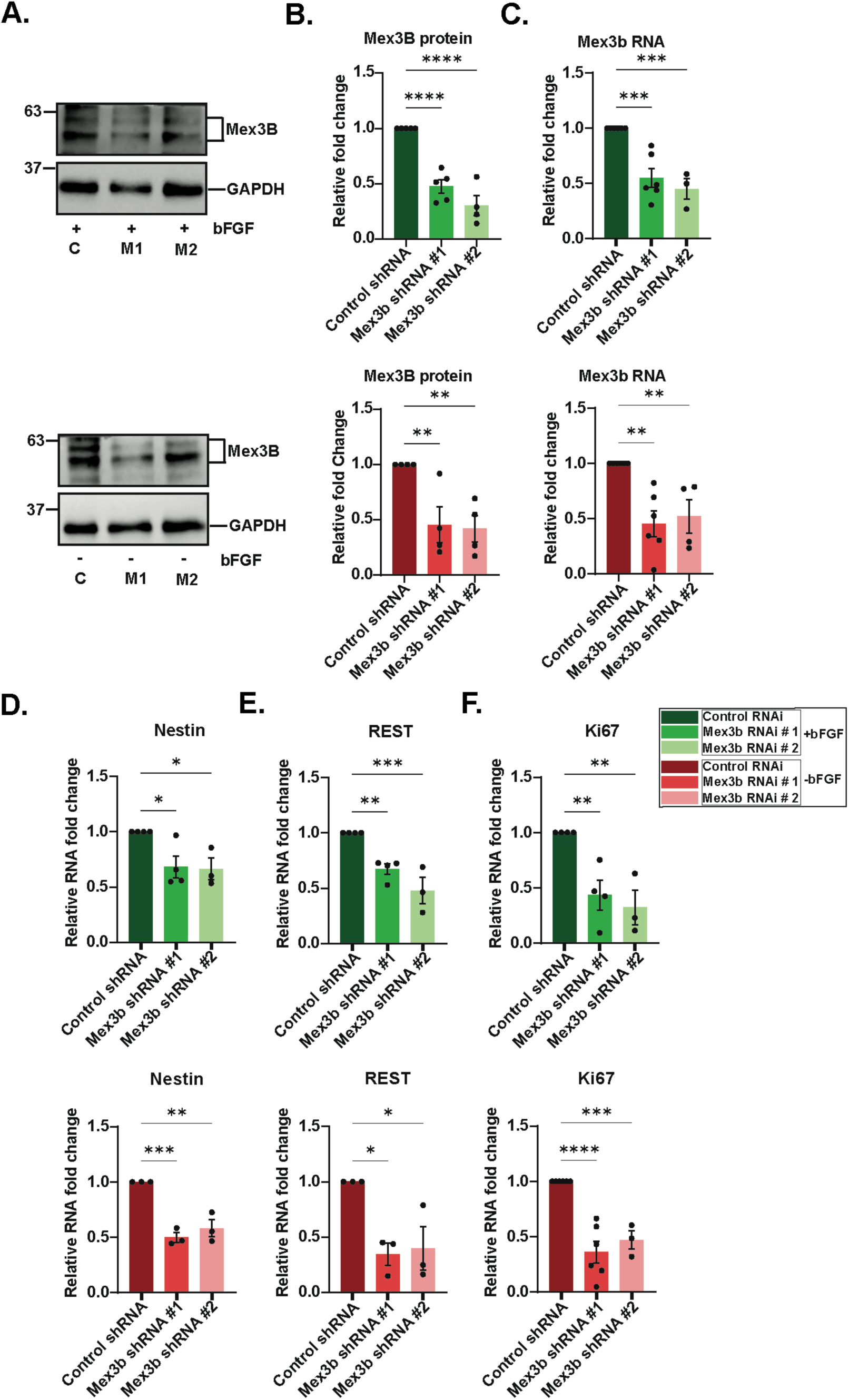
MEX3B reduces the expression of RNA of proliferation markers in NPCs. **A-B.** Western blot quantification of MEX3B knockdown in NPCs in plus and minus-bFGF expressing shRNA as indicated. n=3-5, **p<0.01, ****p<0.0001. (C=Control shRNA, M1=Mex3b shRNA #1, M2=Mex3b shRNA #2) **C.** qRT-PCR based quantification of *Mex3b* RNA in NPCs in plus and minus-bFGF expressing shRNA as indicated. n=3-6, **p<0.01, ****p<0.0001. **D-F.** Differential expression of proliferation markers D-Nestin, E-REST, F-Ki67 in NPCs in plus-bFGF and minus-bFGF on MEX3B knockdown as indicated. n=3-6, *p<0.05, **p<0.01, ***p<0.001, ****p<0.0001. All the data is shown as Mean±SEM and One-way ANOVA with Fisher’s LSD is used for statistics.

### MEX3B protein alters the translation of neural proliferation and differentiation markers in neuronal progenitors

To test the translational expression of proliferation and differentiation markers in NPCs we enumerated the Ki67-positive, NeuN-positive, MAP2-positive, and GFAP-positive cells were immunostained with antibodies against the respective markers. NPCs were grown on coverslips and infected with the lentivirus-containing plasmid expressing shRNA against Mex3b and EF1α-GFP for immunostaining at DIV5-6 (Fig.S4A). The enumerated marker-positive cells were plotted as the percentage with respect to DAPI-positive cells. We observed a significant reduction in the percentage of Ki67 positive cells (Mex3b shRNA construct #1: 11.84±5.659% decrease; p=0.0397; construct #2: 13.59±5.366% decrease; p=0.0133) which is a proliferation marker while the percentage of NeuN positive cells (Mex3b shRNA construct #1: 9.815±4.543% increase; p=0.0339; construct #2: 9.862±4.642% increase; p= 0.0369) and MAP2 positive cells (Mex3b shRNA construct #1: 28.08±8.591% increase; p=0.0016; construct #2: 28.83±8.447% increase; p=0.001) (Fig.4A-F) which are neural differentiation marker increased significantly upon MEX3B knockdown even in the presence of a very strong proliferative push bFGF. Furthermore, no change was observed in the levels of glial marker GFAP (Mex3b shRNA construct #1:0.02743±3.346% higher; p= 0.9935; construct #2: 0.4873±3.346% lower; p=0.885) (Fig.4G-H). Absence of MEX3B in NPCs pushes neural progenitors to attain the neural fate without affecting the glial fate of NPCs by asserting a translational control on specifically neural differentiation markers.

**Figure 4:**
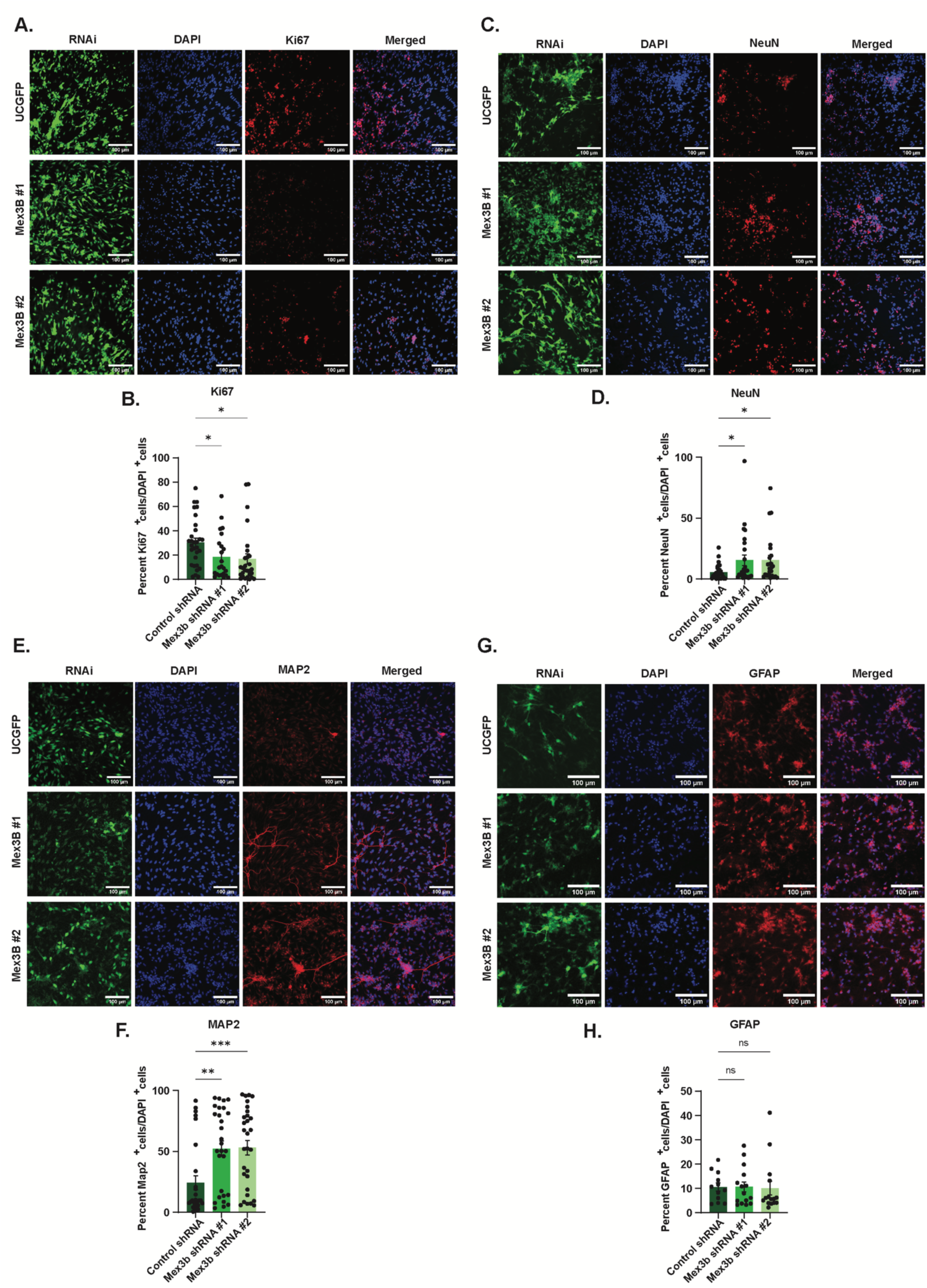
MEX3B knockdown affects proteomic expression of proliferation and differentiation markers in NPCs. **A,C,E,G.** Representative images of NPCs expressing indicated shRNA immune-stained with antibody against Ki67 (A.), NeuN (C.), MAP2 (E.), GFAP (G.) and co-stained with DAPI. **B.** Percentage quantification of Ki67^+^ cells per DAPI^+^ cells represented in A. n=23-30, *p<0.04. **D.** Percentage quantification of NeuN^+^ cells per DAPI^+^ cells represented in C. n=24-29, *p<0.04. **F.** Percentage quantification of MAP2^+^ cells per DAPI^+^ cells represented in E. n=28-30, **p<0.002, ***p<0.001. **H.** Percentage quantification of GFAP^+^ cells per DAPI^+^ cells represented in G. n=12-15, ns=not significant. All the data is shown as Mean±SEM and One-way ANOVA with Fisher’s LSD is used for statistics. Merged=composite image of DAPI and respective immuno-stained image. Scale bar=100µm.

### Knockdown of MEX3B switches the proliferative state of NPCs to a differentiated neuronal fate

We next looked for morphological changes, if any, induced by MEX3B. NPCs being dividing progenitor cells are mostly round to elliptical with no to very few appendages while a differentiated NPC has developing primary to secondary neurites and, axons, etc. We have exploited this property of differentiated NPCs to distinguish them from progenitors and study if MEX3B have structurally effects on the neural progenitor cell fate. The percentage of cells with primary neurites (counted using AutoneuriteJ) to that of DAPI-positive cells was calculated. RNAi with Mex3b shRNA (construct #1: 12.14±2.179% increase; p=<0.0001; construct #2 14.98±2.179% increase; p=<0.0001) showed a significant increase in the number of cells with primary neurites compared to Control shRNA in the NPC grown in the presence of bFGF (Fig.5C-D). Neurite length of the neurites was also found to be increased significantly in MEX3B RNAi (shRNA construct #1: 590±120.6 increase; p=<0.0001; construct #2: 607±122.2 increase; p=<0.0001) compared to the Control RNAi (Fig.5A-B). The Sholl analysis performed also showed significantly increased dendritic complexity between 10um distance from soma to 120um distance from the soma in MEX3B knockdown NPCs compared to Control (Fig.S4B-C). Hence, the results corroborate the negative effect of MEX3B on primary neurite outgrowth and neurite length in differentiating NPCs.

**Figure 5:**
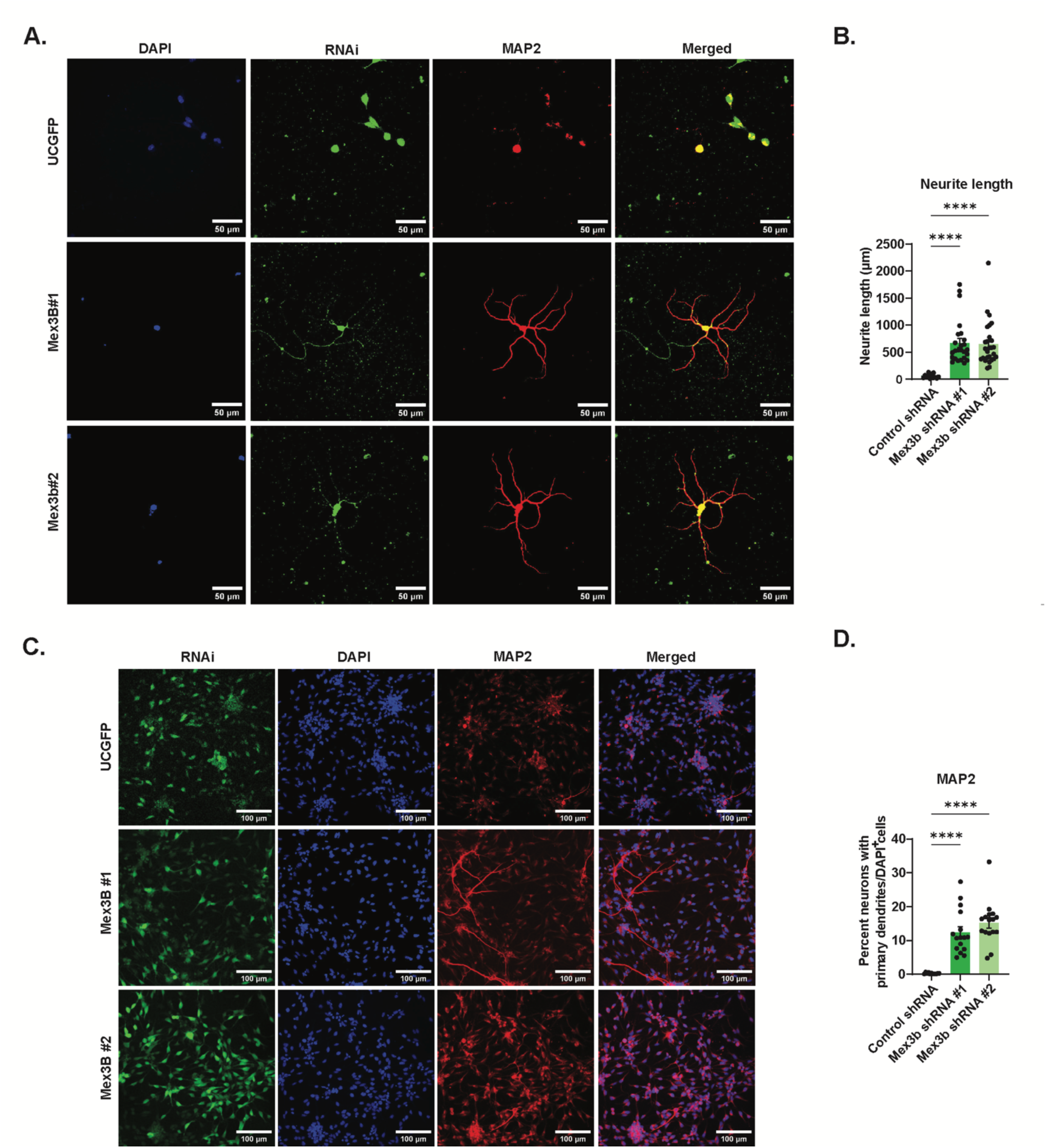
Loss of function of MEX3B promotes neuronal differentiation. **A and C.** Representative images of NPCs expressing indicated shRNA immune-stained with antibody against MAP2 co-stained with DAPI (magnification to 60X for A. Scale bar= 50µm and 10x for B. Scale bar=200µm). **B.** Neurite length measurements in µm of differentiated NPCs represented in A. done using SynD. n=14-25 NPCs, ****p<0.0001. **D.** Quantification of Percentage of cells with primary dendrites per DAPI^+^ cells represented in C. using AutoneuriteJ Image J plugin. n=11-16, ****p<0.0001. All the data is shown as Mean±SEM and One-way ANOVA with Fisher’s LSD is used for statistics.

### MEX3B promotes the proliferation of NPC by associating with REST and HOTAIR

To delve deeper into understanding the role of Mex3b in the proliferation of NPC we looked into the literature for any mechanistic insights on Mex3b. Notably, two molecules emerged as particularly significant to explore further: the long non-coding RNA HOTAIR (HOX Transcript Antisense RNA) and the transcriptional repressor REST (RE1-Silencing Transcription factor). MEX3B has been found to associate with HOTAIR for its ubiquitination activity in HEK293T cells [56]. Since, HOTAIR is known to be involved in developmental defects [57] and brain disorders [58–60], it is pertinent to examine its interaction with MEX3B in proliferating NPCs. REST, on the other hand, is considered as the master switch for neural fate determination [52, 53, 61]. Additionally, our previous experiments have indicated that Mex3b also plays a role in determining neuronal fate and The dataset from Ma’ayan Laboratory of Computational Systems Biology indicated the interaction of REST transcription factor with *Mex3b* mRNA [62]. Moreover, HOTAIR has been reported to interact with REST as part of Polycomb Repressive Complex 2 (PRC2) and LSD1/CoREST/REST complex [63]. Thus, exploring the interplay between MEX3B, HOTAIR and REST in the context of NPC proliferation and differentiation is essential for elucidating the molecular mechanism governing neural development. To begin with, we tested the effect of MEX3B knockdown in NPCs on REST and HOTAIR expression levels. Previously, we have shown reduction in REST transcript on MEX3B knockdown in proliferating and differentiating NPCs (Fig.3E). Now, REST protein levels are also observed reduced several folds (Mex3b shRNA construct #1: 0.4488±0.06399 fold decrease; p=0.0009; construct #2: 0.5559±0.06399 fold decrease; p=<0.0003) upon MEX3B knockdown in proliferating NPCs as well in minus-bFGF condition (Mex3b shRNA construct #1: 0.6016±0.1768 fold decrease; p=0.0078; construct #2: 0.6875±0.1768 fold decrease; p=0.0037) (Fig. 6A and B).

**Figure 6:**
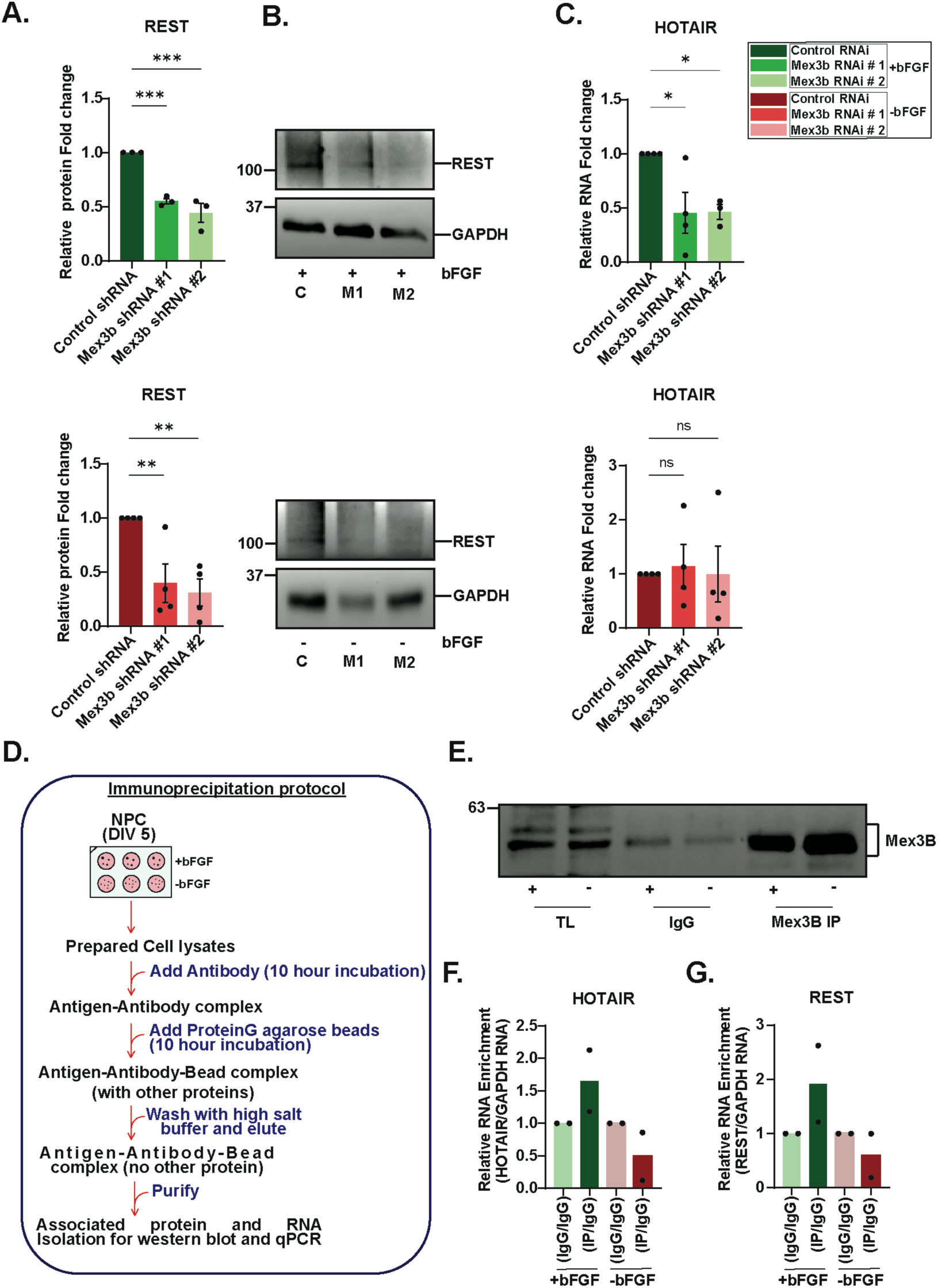
MEX3B protein associates with REST and HOTAIR RNA in proliferating NPCs. **A-B.** Western blot quantification of REST protein levels upon Mex3B knockdown in NPCs in plus and minus-bFGF expressing shRNA as indicated. n=3-4, ***p<0.001, **p<0.01. **C.** qRT-PCR quantification of HOTAIR RNA in NPCs in plus and minus-bFGF expressing shRNA as indicated. n=3-4, *p<0.05, ns=not significant. **D.** Schematics of Immunoprecipitation protocol. **E.** Immunoblot representing enriched expression of MEX3B in pulldown samples(TL= total lysate, IgG and Mex3B IP) in plus and minus-bFGF conditions. **F-G.** Histogram representing enriched expression of HOTAIR (F.) and REST (G.) RNA in Mex3B pull down sample in plus-bFGF condition. All the data is shown as Mean±SEM and One-way ANOVA with Fisher’s LSD is used for statistics.

Similarly, HOTAIR levels were also checked. HOTAIR expression was reduced on MEX3B knockdown in plus-bFGF condition (Mex3b shRNA construct#1: 0.5468±0.1685 fold decrease; p=0.0118; construct#2: 0.5362±0.182 fold decrease; p=0.0185) compared to control shRNA but remain unchanged in minus-bFGF condition (Fig. 6C). The variations observed in REST and HOTAIR expression in our system (Fig. S5A-C) and on MEX3B knockdown and the observations from previous studies prompted us to study the interaction between the three molecules.

To test our hypothesis, we immunoprecipitated MEX3B protein using anti-MEX3B antibody and checked for HOTAIR lncRNA and REST mRNA enrichment in MEX3B-IP samples compared to non-binding control samples in both with and without bFGF condition (Fig. 6D and E). We discovered that both HOTAIR lncRNA (plus-bFGF: 0.6531 fold increase; minus-bFGF: 0.5099 fold decrease) and REST mRNA (plus-bFGF: 0.918 fold increase; minus-bFGF: 1.328 fold decrease) shows strong binding in plus-bFGF condition relative to non-binding control but the binding is weak in minus-bFGF condition (Fig. 6F and G). Stronger association of MEX3B protein with HOTAIR and REST mRNA in proliferating NPCs emphasized that interplay between MEX3B, HOTAIR and REST is integral to the maintenance of proliferative capacity in progenitors.

## Discussion

This study highlighted the role of the E3 ubiquitin ligase MEX3 (Muscle Excess 3) family member, MEX3B, as an RNA-binding protein in the proliferation of neural progenitors. While E3 ligases have been very well characterized in various contexts of neuronal proliferation and development [20–25], what remains to be comprehensively studied is how a subset of E3 ligases, possessing RNA binding domains (known as RNA-binding Ubiquitin ligases or RBULs) influence the post-transcriptional and translational fates of mRNAs involved in the determination of cellular identities. Since, there exists a large-scale dynamicity in the temporal patterns of gene expression during neuronal proliferation and differentiation, RBULs become interesting candidates to study because the dual possession of ubiquitination and RNA binding properties allow RBULs to have control over RNA metabolism as well as protein turnover [64]. The focus of this study was to identify whether RNA binding properties of E3 ligases provide any non-canonical advantages to their influence over the proliferation or differentiation events associated with neuronal development.

We performed microarrays from neural progenitor cell cultures (NPCs), taken from the cortex of E11.5-E12.5 mouse embryos and grown in the presence and absence of bFGF, a pro-proliferative cue. Cortical progenitor cells maintain their proliferative state in the presence of bFGF, its withdrawal results in the onset of differentiation, marked by the formation of neurites. NPCs, therefore, provides a viable environment to investigate the influence of E3 ligases in the coordinated transcriptional and translational paradigms that occur during the transition phase between proliferation and differentiation. Microarray studies from NPCs grown in the presence or absence of bFGF revealed a subset of differentially expressed E3 ligases. Cross-comparison of the list of RNA Binding proteins (RBPs) in mouse [54] with the total number of E3 ligases in our microarray (seventy-six), revealed that 17 of the E3 ligases were RBULs. Further GO analysis (using PANTHER) of the differentially expressed E3 ligases in our microarray revealed that amongst them, the ones that were common to the PANTHER class “RNA metabolism” and “binding” were the MEX3 family of E3 ligases (Figure 1). Single qRT-PCR of the MEX3 family candidates confirmed that *Mex3B* mRNA displays maximum change between proliferative neuronal progenitors and their differentiated forms. (Figure 2 and S2). MEX3B’s function, as an E3 ligase through its RING finger domain [51] is well known but its potential as an RNA-binding protein remains elusive. Mex3, a homolog of the Mex3 family in *C. elegans* is known to be involved in the germline totipotency [65] and very recently the role of Mex3B in the spatial organization of Sertoli cells to maintain spermatogenesis in testis in mouse has been identified [66]. A homolog of Mex3a has been reported as the regulator of nervous system development in a vertebrate *Xenopus laevis* [67, 68]. However, little to none is known about *Mex3b* expression in the context of neuronal proliferation in vertebrates. From the Mouse ENCODE transcriptome data, *Mex3b* transcript levels were found to be specifically higher in the whole brain at E14.5 and in CNS at E11.5, E14, and E18 (Embryonic day) [69]. Differential expression of *Mex3b* mRNA in microarray data further strengthened our hypothesis that Mex3b is involved with the proliferation/differentiation of neural progenitors (Figures 1 and S1).

Since MEX3B is an E3 ligase and an RNA-binding protein, we needed to assess whether it would be functionally active during neuronal proliferation. The expression of MEX3B protein was surprisingly found to be significantly high in +bFGF condition compared to -bFGF condition which is completely contrasting to *Mex3b* mRNA levels. Previous studies helped us understand this ambiguity. A study from the Masanari Taira’s group shows the autoregulation of *Mex3b* mRNA by its protein in *Xenopus levis*. MEX3B binds to the 3’UTR of its own mRNA to destabilize it by regulating the post-transcriptional expression, hence the protein and RNA have opposite expression patterns at distinct temporal periods during development [55]. Although not E3 ligases, several other proteins regulate their mRNA; for example Mel-N1 in mice increases the stability of its mRNA by competing with destabilizing factors for its binding site on the 3’UTR region [70–72]. Another protein TTP shares considerable similarity in regulation of its own mRNA with MEX3B through binding to AUUUA sequences [73]. We enquired whether such autoregulation of *Mex3b* mRNA by its protein is valid in vertebrates as well. Accordingly, we assessed Mex3b protein and mRNA levels across different time points, both *in vivo* and *in* vitro. Within NPCs, Mex3b protein and mRNA expression patterns were followed in both +bFGF and –bFGF conditions. Interestingly, in the presence of bFGF, *Mex3b* mRNA displays a steep decline in expression levels with time, with a concomitant increase in MEX3B protein levels (Figure 2D-2G). This is similar to the auto-regulatory effect of MEX3B protein on its mRNA as reported in *Xenopus* [52]. Our observation bolsters the idea that increased expression levels of MEX3B protein facilitate proliferation over differentiation in NPCs. The autoregulation of *Mex3b* mRNA by its protein is also dependent on the presence of bFGF; we infer this from the expression of Mex3b protein and mRNA in NPCs that have ceased to proliferate in the absence of bFGF. Indeed, in –bFGF conditions, there is no significant difference in protein and RNA levels of *Mex3b*, suggesting that the presence of bFGF is imperative for the contrasting expression pattern of Mex3b protein and it’s mRNA (Figure 2). Physiologically, the expression of MEX3B protein in the cortex of CD1 mouse was significantly low at E11.5, E15, and P0, but mRNA levels were high, Our data corroborates with the high levels of *Mex3b* transcript reported at similar developmental time points in Mouse ENCODE transcriptome data [69]. The discrepancy in the expression pattern of MEX3B protein in NPCs as compared to that in mouse cortex can be attributed to the simultaneous influence of multiple growth factors, in different areas of the developing cortex. Considering that the vertebrate cortex has six layers, it is highly probable that the synergistic influence of various pro-proliferation and pro-differentiation cues during cortical development result in the observed pattern of Mex3b expression. In contrast, the sole proliferative cue provided in NPCs is bFGF, therefore the observed increase in MEX3B protein levels during proliferation can be attributed to bFGF’s overwhelming influence. The steep increase of MEX3B protein levels in mouse cortex at P7 and P14 can be an interesting observation to explore further. Certain parts of the cortex develop during this time window, including important developmental events such as eye and ear opening [74].

We performed MEX3B RNAi within NPCs to understand its influence on various aspects of neuronal proliferation and differentiation. A neural progenitor cell with proliferative property can be well distinguished from a differentiated neuron by measuring different parameters like the differential expression of proliferative and differentiation markers, and the morphological differences observed in progenitor cells and differentiating cells. Transcriptional expression of proliferative markers like Ki67, Nestin, and REST reduced significantly upon MEX3B knockdown even in the presence of bFGF; indicating that the absence of MEX3B pushes the fate of neural progenitors towards differentiating cells; circumventing the influence of bFGF (Figure 3 and S3). Ki67 is a generic proliferative marker and not neuron-specific, but REST and Nestin are specific to neural progenitor proliferation. REST is considered the master switch for the fate determination of neural progenitors [53, 61]. REST protein blocks the transcription of differentiation markers to maintain the proliferative state of the neural progenitor [52, 61]. No change in the transcriptional expression of differentiation markers was observed (Figure S3). Therefore it stands to reason that unlike REST, MEX3B does not influence the transcriptional expression of differentiation markers. However, the translation of both proliferation and differentiation markers is affected. Immunocytochemistry data shows that upon Mex3B knockdown, the number of cells expressing Ki67 protein goes down while the number of cells expressing MAP2 and NeuN increases. MAP2 and NeuN are proteins expressed in fully differentiated mature neurons (Figure 4) [75, 76]. The affected protein levels and unaffected transcript levels of differentiation markers point towards the post-transcriptional control of these markers by MEX3B (Figure 4). MEX3B RNAi resulted in a significant increase in both the percentage of cells with primary neurites as well as neurite length which are the indicators of the greater number of differentiated cells in the pools of progenitors in +bFGF condition. Along with neurite length, changes were also observed in dendritic arborization (Figure 5 and S4). The cumulative observations conclude that the pro-proliferative push of MEX3B is not just molecular but also morphological.

Now, several hypotheses can be posed on how Mex3b is influencing neural progenitor fate. We saw a significant change in both REST mRNA and protein upon MEX3B knockdown (Figure 6 and S5); which led us to further investigate the RNA-binding behaviour of Mex3B. We immunoprecipitated Mex3B protein and assessed the association of *REST* mRNA and the lncRNA *HOTAIR*. MEX3B binding to *REST* mRNA and *HOTAIR* was found to be higher in the +bFGF condition than -bFGF condition. *HOTAIR* is previously been reported to interact with Mex3B and other E3 ligases to facilitate the ubiquitin-mediated degradation of its Snurportin [56]. Mex3B has also been reported to ubiquitinate other targets like Runx3 again in association with *HOTAIR* [77]. *HOTAIR* interacts with REST resulting in transcriptional inactivation or gene silencing [63]. Furthermore, a database released by the Ma’ayan Laboratory of Computational Systems Biology shows target genes of the REST transcription factor in low- or high-throughput transcription factor functional studies from the CHEA Transcription Factor Targets dataset. *Mex3b* mRNA is one of the targets mentioned among 4675 other targets [62].Based on our observations and previous literature, we hypothesize that MEX3B protein acts in concert with *HOTAIR* and REST to regulate proliferation in NPCs. The stability of REST mRNA may also be dependent on MEX3B, since knockdown of the latter affects REST mRNA and protein levels (Figure 3 and Figure 6). MEX3B is reported to share molecular functional similarities with PUF-8 in regulating mitosis/meiosis in C.elegans [78–80]. So plausibly MEX3B has the potential to be an RNA-binding protein involved in *REST* mRNA regulation through its 3’UTR. MEX3B has been reported to provide resistance to cancer immunotherapy by downregulating the expression of *HLA-A* mRNA [81, 82].

Our study opens a new avenue for the transcriptional/post-transcriptional regulation of neural fate determination through the transcriptional control of REST by an (RBUL) RNA binding ubiquitin ligase protein MEX3B. It also provides a link between an lncRNA, an RUBL protein, and the transcription factor REST and neuronal differentiation. We explored the non-canonical function of Mex3b and tried to address the complex interplay between the RNAs RBPs; and how it may dictate cellular phenotypes. The physiological impact of this intricate network of interactions on neural proliferation and differentiation is also answered through our study. Henceforth, this study along with more such studies in future can help us to understand and treat certain neurodevelopmental disorders caused due to defects in neuronal proliferation and differentiation.

## Conclusion

MEX3B belongs to a special class of E3 Ubiquitin ligases that also possess RNA binding motifs. Our study provides an in-depth characterization of MEX3B’s influence on the proliferative properties of neuronal progenitor cells (NPCs). Structural changes in proliferating NPCs, attainment of neuronal fate and the display of characteristics similar to differentiated neurons upon MEX3B RNAi further underline its involvement in neural fate determination. From the observations obtained from our study we can conclude that MEX3B brings about a strong pro-proliferative drive that aids in the occlusion of differentiation programme in NPCs. Mechanistically, MEX3B associates with the master switch REST along with an lncRNA *HOTAIR*, known for its role in neuronal development. Overall, the study emphasizes that MEX3B-REST-*HOTAIR* axis may be integral to a complex cascade of events that maintain neural progenitors in a proliferating phase. Our study provides a hitherto uncharacterized example of an E3 ligase using its RNA-binding property to spatiotemporally regulate an intricate molecular network underlying neural development.

## Material and Methods

### Animal

Time-pregnant female CD1 mice were used for the cultures. Animals were issued from the in-house National Brain Research Centre Animal house facility. Animal issuing protocol numbers NBRC/IAEC/2023/184, are approved by IAEC (Institutional Animal Ethics Committee) of NBRC registered with the CPCSEA (Committee for the Purpose of Control and Supervision of Experiments on Animals) (Registration number: 464/GO/ReBi-S/Re-L/01/CPCSEA). Animals were maintained well at room temperature (25± 0.5 °C), humidity 40 ± 5% with a 12-hour light/dark cycle along with food and water *ad libitum*. Mating pairs were set carefully as per requirements. Regular examination was conducted for the positive vaginal plug. Embryonic day 0 (E0) was noted as the date of the plug. Pregnant animals were maintained and checked for weight till E11.5 the date of culture.

### NPC (Neural Progenitor Culture)

100mm dishes were coated with PDL (50ug/ml) and Laminin (5ug/ml) incubated overnight. Washed with autoclaved milliQ water and kept for drying. 7.5ml (Neurobasal (#21103049) supplemented with B27 (#17504044)) NB27 plus bFGF (40ng/ml) media was added to these 100mm dishes.

Dissected out the pups from the E11.5-12.5 time pregnant mother in to dissection media, taken out the forebrain kept in NB-27 plus-bFGF media. Washed twice with NB-27 plus-bFGF media. Triturated gently in 1ml NB-27 plus-bFGF in 1.5 ml microcentrifuge tube to obtain 4 to 8-celled clumps. Observed the neurospheres under the microscope. Added 500ul of neurosphere/cell suspension to the tube containing 4ml fresh NB-27 plus-bFGF. Plated this onto the two 100mm dishes. Maintained the culture till it reached confluency by changing half media every alternate day. On day7 Neurospheres were dissociated using 0.25% trypsin, DMEM with 10%FBS was added and cells are centrifuged at 800rpm at room temperature to remove trypsin. Resuspended in fresh NB-27 plus-bFGF media. Visualized and counted the cells using a haemocytometer and plated 150000 cells per well on 6 well plates. Initially, all cells were plated in NB-27 plus-bFGF media, after 8 hours the media for minus-bFGF wells was changed to NB-27 without bFGF.

### Microarray data and GO analysis

NPC cultures with bFGF and without bFGF were grown as described above for microarray analysis. Antisense against all the E3 ligases was used to check the differential expression of these E3 ligases in the two conditions. The microarray data obtained was normalized with GAPDH and heatmap was made using GraphPad Prism9.

Gene ontology analysis was performed for all the E3 ligases used in the microarray data and for the Mex3B targets obtained from previous studies using the PANTHER Database (http://pantherdb.org/) (doi:10.5281/zenodo.10536401). Panther statistical overrepresentation analysis was performed with a statistical test type chosen to be Fisher’s Exact and corrected for False discovery rate. Fold Enrichment of the transcripts in annotated datasets such as GO: Biological Process and GO: Cellular Components, GO: Molecular function was generated along with FDR values. Significantly enriched GO terms were analyzed and plotted in GraphPad Prism9. Venn diagram was made using online available tools for finding common proteins in the RNA binding proteins reported so far and the E3 Ubiquitin ligases used in our microarray data.

### Lentivirus preparation

shRNA against Mex3B shown in Table 1:

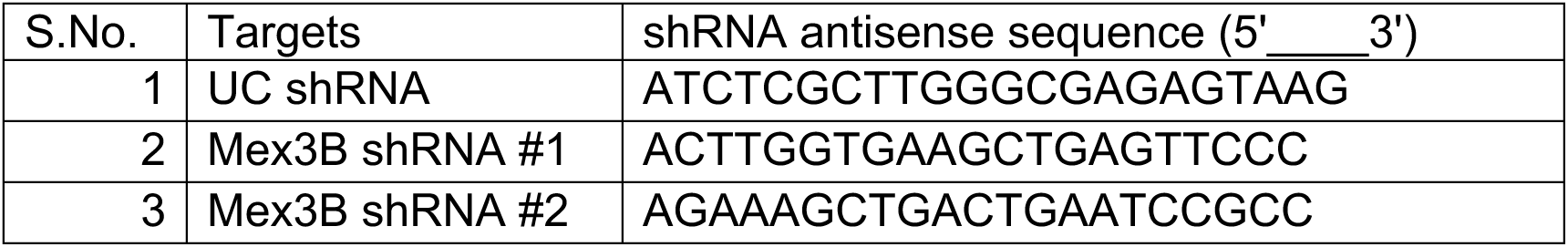

shRNAs were cloned into the pLVTHM plasmid (Addgene #12247) at MluI and ClaI sites. The cloned vectors were verified by sequencing. The Escherichia coli Stbl3 strain was used for the propagation of pLVTHM plasmids containing shRNA, while DH5 alpha was employed for propagating the psPAX2 (Addgene plasmid #12260) packaging plasmid and the pMD2.G (Addgene plasmid #12259) envelope plasmid. Purified plasmids were obtained using the Endotoxin-free Maxiprep kit (Qiagen). Lentiviruses were generated by co-transfecting 20 mg of the transfer vector (mCherry cassette under the EF1alpha promoter and shRNA cassette against Mex3B or non-targeting control under the H1 promoter in the pLVTHM plasmid), 15 mg of psPAX2, and 6 mg of pMD2.G into HEK293T cells.

HEK293T cells were cultured in low-glucose DMEM media (Gibco) with 10% Fetal Bovine Serum (FBS) (Gibco) at 5% CO2 and 37°C. Transfection of HEK293T cells (2 x 106) was performed using the calcium phosphate method. Following transfection, the media containing the transfection mixture was replaced with fresh media after 8 hours. Culture media containing lentivirus particles, was collected 72 hours post-transfection, and the viruses were concentrated through ultracentrifugation and resuspended in NB-27 media. Viral titers were determined by infecting HEK293T cells and counting the cells using haemocytometer, with typical titers ranging from 1 to 3 x 107 TU/ml.

RNA interference (RNAi) was carried out by infecting neural progenitor cells (NPCs) at Days-In Vitro (DIV) 0, which is 8 hours post-plating, with lentivirus expressing shRNAs targeting Mex3B and the non-targeting control. 50% of media was changed every alternate day. Virus infection was tracked by observing GFP expression in infected cells.

### Knockdown efficiency check

The efficiency of the knockdown was confirmed by isolating RNA and protein from Mex3B and non-targeting shRNA virus-infected cultures at DIV 5 using Trizol and 1X RIPA buffer respectively and further processing the samples as mentioned below.

### RNA isolation and cDNA preparation (from NPC and from animals)

At day 5 post-NPC culture, samples from 3-6 wells of a 6-well plate were pooled to isolate RNA from equal number of cells for each experimental condition. Cells were washed with 1xPBS to remove media completely and RNA isolation was done using Trizol reagent. Samples were DNase treated using (Invitrogen). cDNA was prepared using an Invitrogen Superscript-III kit and qPCR was performed using SYBR green (Applied Biosystems).

For the physiological relevance of Mex3B. CD1 mouse at different developmental time points viz. (embryonic days) E11.5, E15, (Postnatal days) P0, P7, P14, P21, P28 was sacrificed and isolated cortex. Homogenized the cortex in Trizol to isolate RNA. cDNA was prepared using the Invitrogen kit and qPCR was performed using SYBR green.

### Timepoint NPC sample collection

For time point experiments samples from 3-6 wells of NPCs grown in 6 well plates were collected at 1 hour, 3 days, 5 days, 7 days from the time point of changing the media from NB-27 with bFGF to NB-27 without bFGF. Cells were washed with 1xPBS to remove media completely and collected in a Trizol reagent. RNA and protein both were isolated from the same Trizol collected cells. The aqueous phase was used for RNA isolation and remaining phase was used for protein isolation using a predefined protocol. RNA was used for qPCR and protein for western blot.

#### qPCR primers

All qRT-PCR primers have been listed in Table 2.

**Table 2:**
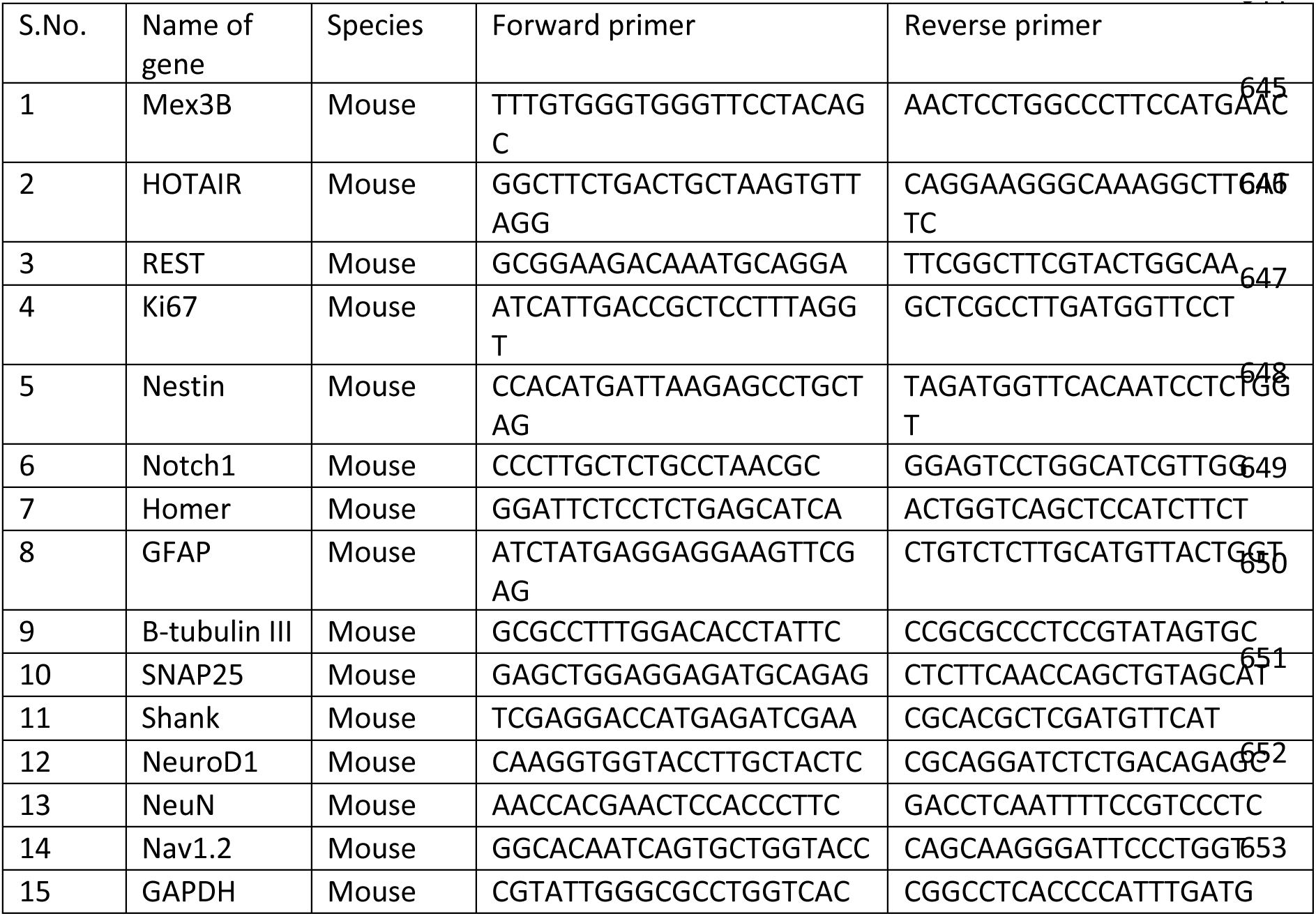

### Protein Isolation and Western blot (from NPC and from Animal)

We collected samples from various experimental conditions and lentivirus-infected neuronal progenitor cultures, each originating from two wells of 6-well plates. These samples were lysed using 1X RIPA buffer for western blot analysis. After quantitatively estimating the protein concentration in each sample, equal amount of lysates was loaded onto 8-10% SDS polyacrylamide gels. The resolved proteins were then transferred onto nitrocellulose membranes (Millipore), followed by blocking with 5% BSA (Merck).

The blots were probed with specific antibodies: Mex3B (1:500; ThermoFisher, #A301-194A), Tuj1 (1:10,000; Sigma, #T8578), GAPDH (1:10,000; Sigma, #G9545) and REST (1:500; Millipore, #09-019). For each lane, immunoblotting was performed using GAPDH as the internal control to normalize protein levels. To assess the developmental expression pattern of Mex3B, we analysed cortical tissue lysates obtained from various developmental stages, including embryonic day (E11.5, E15) and postnatal day (P0, P7, P14, P21, and P28), via SDS-PAGE with Mex3B antibody. GAPDH expression was utilized for normalization purposes. Protein estimation was carried out using BCA protein estimation kit (Invitrogen).

### Immunocytochemistry

For Immunocytochemistry 20 thousand NPCs were plated in the 12 well plates with 18mm coverslips coated with poly-d-lysine and laminin. 8 hours post plating cells were infected with the virus for Control RNAi and 2 constructs of Mex3B RNAi. At DIV6 the coverslips were fixed with 2% PFA and 4% sucrose solution and subjected to immunostaining, following a previously described protocol [83]with minor adjustments.

NPCs were washed three times with Phosphate Buffer Saline (PBS) and then fixed with 4% paraformaldehyde (PFA) in 1X-PBS containing 4% sucrose at 37°C for 30 minutes. Subsequent to fixation, coverslips were subjected to 3 times 1XPBS washes again, permeabilized with 0.5% Triton-X-100 in 1X-PBS (PBST) for 15 minutes, and then washed with 1XPBS. Blocking was done for 4 hours at room temperature in a blocking solution consisting of 5% bovine serum albumin (BSA) and 0.1% Triton-X-100 in 1XPBS. The NPCs were subsequently incubated with specific antibodies against MAP2 (1:300, Mouse, clone K28/43, Neuromab, #75-028), NeuN (1:500, Goat, Invitrogen, #PA5-143586), Ki67(1:50, Mouse, abcam, ab279653) and GFAP(1:1000, Mouse, Millipore, MAB360) overnight at 4°C. After the primary antibody incubation, NPCs were washed thrice with 1XPBST (0.01% Triton-X-100) and incubated with the respective secondary antibodies (mouse or rabbit or goat) conjugated with Ms-Alexa–546 (Invitrogen Molecular probes, #A-11003) or Ms-Alexa– 633 (Invitrogen Molecular probes, #A-21052) or Rb-Alexa–633 (Invitrogen Molecular probes, #A-21071) or Goat-Alexa-633 (Invitrogen Molecular probes, #A-21082) dyes at room temperature for 2 hour. Following another three washes with 1XPBS (0.01% Triton-X-100), coverslips were air-dried and mounted on glass slides using Vectashield mounting medium containing DAPI (Vector Laboratory, #H1200) and sealed with nail paint.

### Confocal microscopy

Using a Nikon A1HD point scan confocal microscope equipped with a Plan Apo 10X objective, we captured a large image of dimension 11X11 frames, 16-bit images of the entire coverslip containing immunostained NPCs with each image at a resolution of 1024 x 1024 pixels, and an optical zoom of 0.7. We acquired 6-12 optical sections with a step size of 3µm. To facilitate further analysis, the entire coverslip image was divided into multiple non-overlapping equal (Region of Interest) ROIs.

Fluorescence labeling for DAPI, GFP (Lentivirus RNAi), Alexa-546 (GFAP, Ki67), and Alexa-633 (MAP2, NeuN) was achieved by exciting with 405-nm, 488-nm argon ion laser, and 561nm, 640-nm Helium-Neon (HeNe2) laser, respectively. For representative images, Plan Apo 40X objective was used for imaging by Nikon AX point scan confocal microscope. For sholl analysis, images were taken with a Nikon Plan Apochromat 60X with an optical zoom of 0.7 and at a resolution of 1024 x 1024 pixels. We acquired 6-12 optical sections with a step size of 0.75 µm. Consistent laser power, detector gain, amplifier offset, and pinhole diameter settings were maintained for all experiments.

### Cell counting analysis

Cell counts for MAP2 were performed by an experimenter using the ImageJ Plugin Colocalization Object Counter [84]. This plugin allowed the user to establish an auto-threshold for positive labelling in each channel, enabling the identification of cells with DAPI nuclear staining, and cells with the dendritic marker MAP2. For the neuronal dendritic marker MAP2, we determined the lowest noise tolerance that would not erroneously identify a cell as MAP2 positive and considered it as background across our images. Auto threshold was applied consistently across all experimental conditions. In the DAPI channels, the auto threshold was set to include all cells displaying any positive fluorescence. The number of DAPI-positive cells and number of MAP2-positive cells were determined in the respective channels for specific ROI. The data obtained was represented as the percentage of MAP2-positive cells with respect to DAPI-positive cells.

ImageJ Plugin Analyze Particles was used for counting Ki67, NeuN, and GFAP-positive cells. The plugin requires 8-bit images, so the image was processed accordingly by making maximum intensity projections, adjusting the threshold, and converting the image to 8-bit within ImageJ. The best autothreshold type was determined and set for each channel using the autothreshold Plugin in ImageJ for Ki67, NeuN, and GFAP. Similarly, corresponding DAPI images were prepared. ImageJ plugin watershed was used to segregate all the cells. To count the number of cells Particles greater than 10 pixels^2 size was considered as one cell. Circularity parameters were set between 0-0.1. The number of DAPI-positive cells and the number of Ki67/NeuN/GFAP-positive cells were determined in the respective channels for specific ROI. The data obtained was represented as the percentage of Ki67/NeuN/GFAP positive cells with respect to DAPI positive cells.

Statistical significance was assessed using a One-way ANOVA test with Fischer’s LSD post hoc test. All analyses were conducted blindly to the experimental conditions.

### Count of Cells with Primary neurites

ImageJ Plugin AutoneuriteJ [85] was used to count the number of MAP2-positive cells with primary neurites. The plugin creates a masked image of cells with primary dendrites which are also colocalized with the DAPI after thresholding both the channels. Autothreshold chosen to set the threshold for both MAP2 and DAPI channels. Maximum intensity projections were created separately for both channels. Part I of the plugin was used to create the masked images based on the diameter of the nucleus for DAPI staining and the minimal area of neuron provided. Part II of the plugin will count the number of neurons based on the specific neurite length parameter and count the number of joint neurons. The total number of cells with primary neurites is calculated by adding the number of neurons and the number of joint neurons. The data obtained was represented as the percentage of cells with primary neurites with respect to DAPI-positive cells.

### Sholl analysis and Neurite length measurement

We conducted Sholl analysis and neurite length measurements on neural progenitor cells (NPCs) that had been infected with lentivirus and immunolabeled with MAP2 antibody, which served as a dendritic marker for assessing neuronal morphology following RNA interference (RNAi). For this analysis, we employed SynD, a software tool developed by [86]. In this analysis, we first identified the cell soma and proceeded to count intersections within concentric circles, each with increasing radii of 10 µm, originating from the centre of the cell soma. We then plotted the number of intersections against their respective distances from the centre of the cell soma. Total neurite length for each neuron was also obtained from same analysis. To assess statistical significance, we compared the results between the non-targeting control RNAi and Mex3B RNAi samples. To do so, we used Two-way ANOVA as our chosen statistical methods for sholl analysis.

### Immunoprecipitation

Mex3B was immunoprecipitated from NPC cells grown in plus-bFGF and minus-bFGF for 5 days following a previous protocol [87] with minor modifications. On DIV 5 approx. 1.5 million cells (3-6 wells of 6 well plate) were used for IP, media was removed and cells were washed with 1XPBS were collected in Immunoprecipitation buffer (20 mM Tris (pH 7.5), 100 mM KCl, 5 mM MgCl2, 0.5% Nonidet P-40, 1mM DTT, 0.5ug/ul Heparin, 100unit/ml Superase In RNase Inhibitor, 1ul/ml EDTA free Roche Protease inhibitor cocktail) by scraping off all cells using cell scraper. Homogenized using 1ml pipette and mild vortexing. The homogenized lysate was centrifuged at 2000g for 10 minutes at 4degree Celsius, supernatant was collected in a fresh tube for further processing to obtain homogenous lysate without debris. Protein estimation of lysate using a BCA assay kit (Thermo Scientific #23227) was done to use an equal amount of protein for Immuno pull-down. Protein-G agarose beads were equilibrated with immunoprecipitation buffer (20 mM Tris (pH 7.5), 100 mM KCl, 5 mM MgCl2) twice for 5 minutes each. Lysate was precleared with these equilibrated beads for 2 hours, mixing at 50rpm at 4 degrees Celsius. To remove the beads suspension was centrifuged at 8000g, at 4degree Celsius, for 10 minutes. Precleared lysate was divided into two halves one half was incubated with Mex3B antibody and the other half with IgG (non-binding control) with continuous mixing at 50rpm at 4 degrees Celsius for 10 hours. Equilibrated Protein-G-agarose beads were added antibody lysate mixture and incubated with continuous mixing at 50rpm at 4 degree Celsius for 10 hours. Beads with antibody and substrate were collected by centrifugation at 8000g for 10 minutes at 4 degrees Celsius. This matrix of beads was washed once with Immunoprecipitation buffer and then with high salt buffer (20 mM Tris (pH 7.5), 200 mM KCl, 5 mM MgCl2, 0.5% Nonidet P-40, 1mM DTT, 0.5ug/ul Heparin, 100unit/ml Superase In RNase Inhibitor, 1ul/ml EDTA free Roche Protease inhibitor cocktail) 3for 5 minutes and again washed with Immunoprecipitation buffer. Centrifuged at 8000g for 10 minutes at 4 degrees Celsius to collect the washed beads. The pellets were subjected to RNA isolation using the Trizol method with PureLink RNA Mini Kit (Invitrogen), and cDNA was prepared for qPCR. Part of the pellet was kept for protein isolation by boiling in Laemmli buffer to visualize and quantify the result by western blot. Clean blot (Thermo Scientific #21232) was used as a secondary antibody for the western.

### Statistical Analysis

All the data is represented as Mean±SEM. Statistical significance was assessed using the Unpaired two-tailed t-test and corrected for unequal variance by Welch’s correction or by One-way ANOVA test with Fischer’s LSD post hoc test for all qRT-PCR, western blot, and imaging data as indicated with the individual graphs. A two-way ANOVA test with Fisher’s LSD post hoc analysis along with multiple comparisons was used for sholl and NPC time point data as indicated. ‘n’ represents the number of independent experiments conducted to obtain the represented data mentioned along for each experiment. Representative photomicrographs and Immunoblots for each ICC data were prepared using ImageJ. For GO analysis Fisher’s Exact test was used with FDR (False Discovery Rates) value calculations. All the data is analyzed and plotted using the GraphPad Prism 9. GO analysis statistical values were obtained from PantherGO and plotted in the GraphPad Prism 9. The significance cutoff was set at P<0.05 for all the experiments. The respective P values are mentioned with figure legends.

## Acknowledgements

We thank Addgene for Lentiviral vectors. We thank Ishanee Bandyopadhyay for technical assistance. All schematics were made using Biorender.

## Funding

This study was funded by NBRC’s institutional core fund to S.B.

## Author’s Contribution

K.G. and S.S. conceptualized the project and wrote original manuscript. K.G. and G.S. performed experiments and data analysis. K.G., S.S., G.S. critically discussed the experiments and results. S.B. provided funding and initial idea and critical comments on the data and manuscript. K.G., S.S., S.B. edited the manuscript.

## Notes

### Competing Interest Statement

The authors have declared no competing interest.

