## Supplementary Information for "The RNA binding ubiquitination ligase MEX3B regulates bFGF-dependent neuronal proliferation"

**Table S1:** 17 RBULs found in Fig.1E

| Names | Total | Gene Name |
|---|---|---|
| E3 ligases and RNA binding protein | 17 | Cnot4 |
| | | Cpsf1 |
| | | Hectd1 |
| | | Huwe1 |
| | | Mex3a |
| | | Mex3b |
| | | Mex3c |
| | | Mex3d |
| | | Mkrn2 |
| | | Nedd4 |
| | | Nfx1 |
| | | Nfxl1 |
| | | Nosip |
| | | Prpf19 |
| | | Rfwd2 |
| | | Trim2 |
| | | Trip12 |

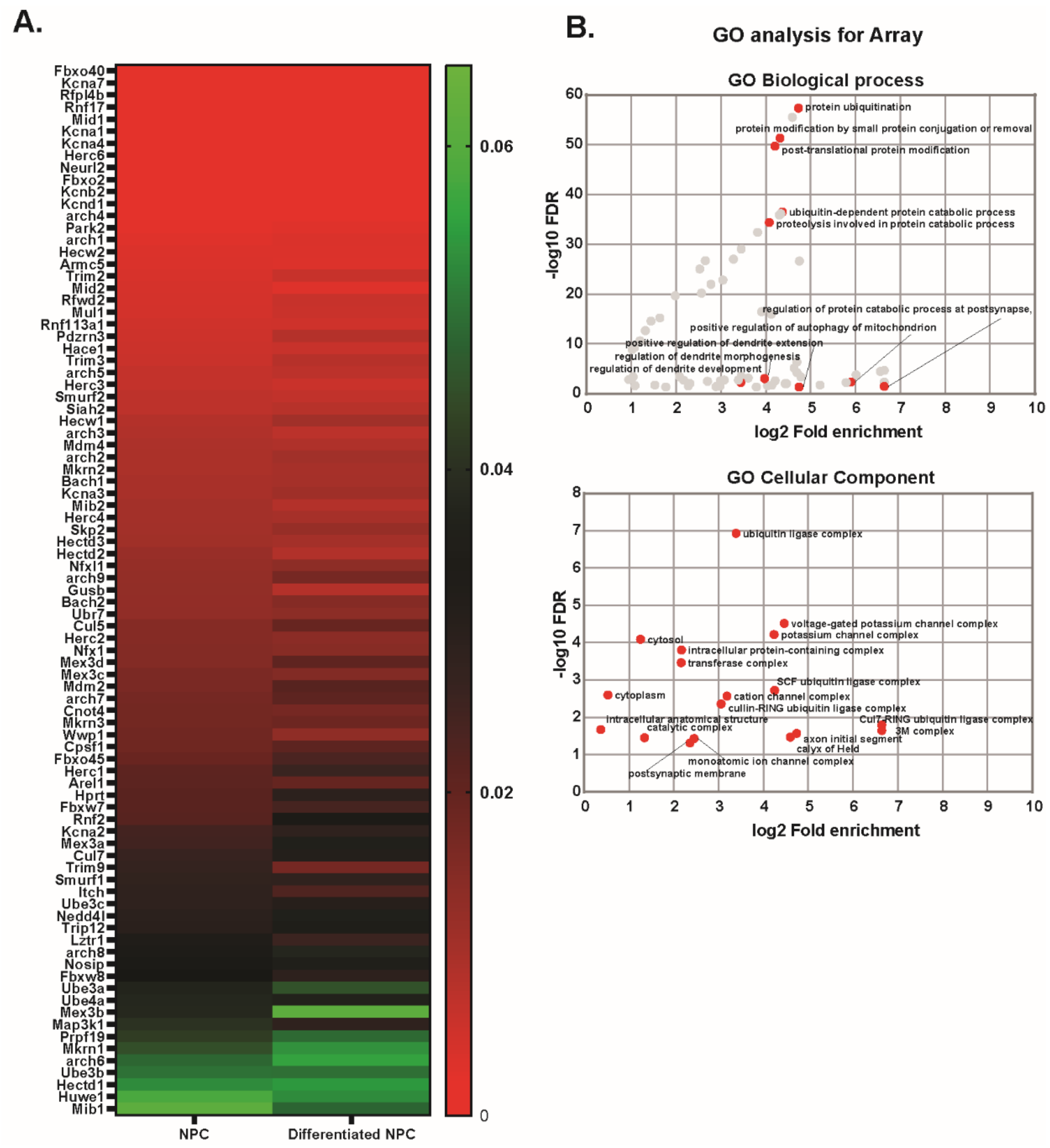


**FigureS1: Delineating the E3 ubiquitin ligase relevant to our system. A.** Heatmap representation of expression of a microarray for E3 ubiquitin ligases in NPCs from n=2. Organized in order of decreasing expression in NPCs. **B.** GO analysis for E3 ubiquitin ligases represented in A.

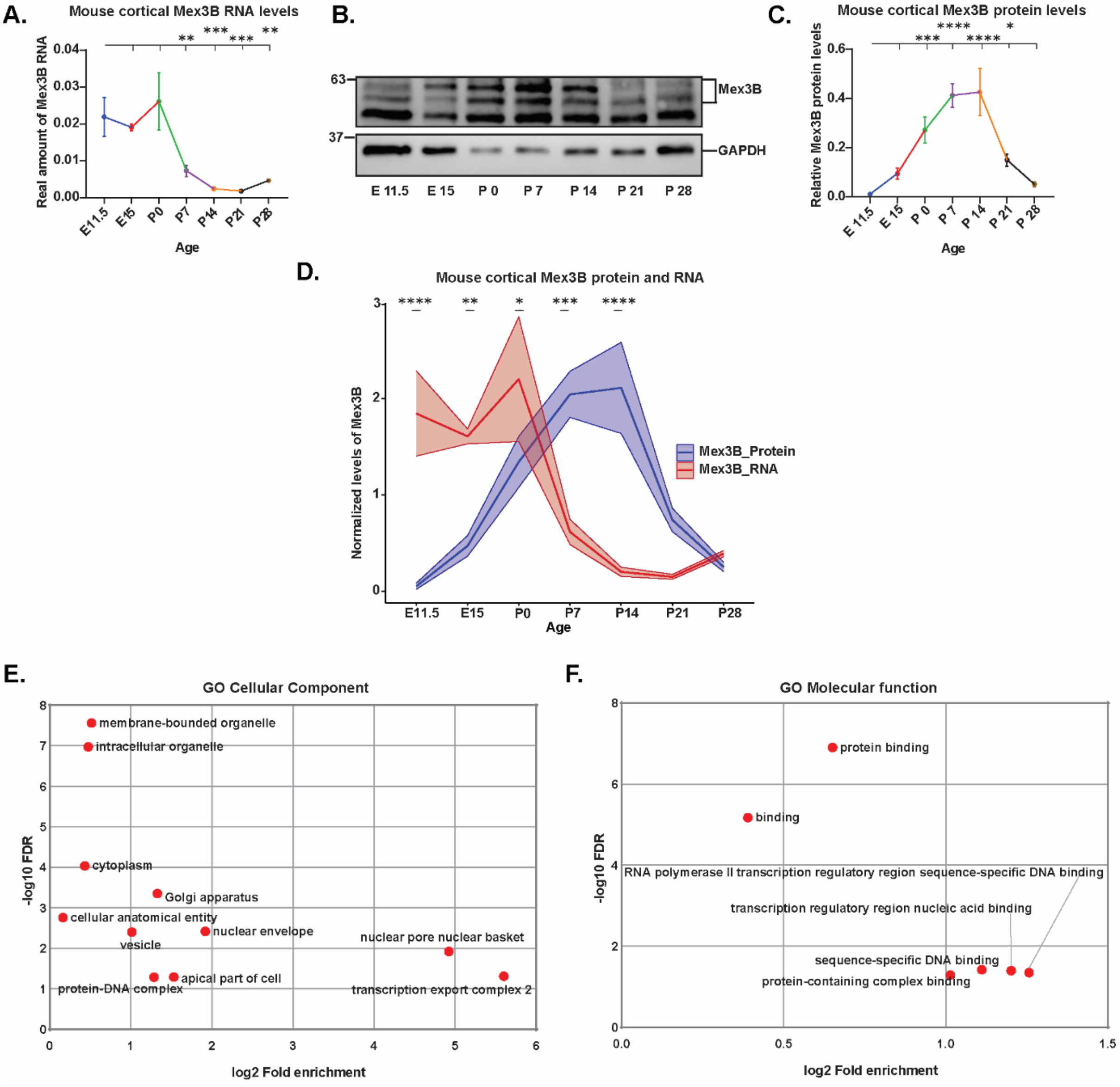


**FigureS2: Inverse correlation between Mex3b RNA and Protein expression. A.** qRT-PCR based quantification of *Mex3b* RNA expression levels in mouse cortex at different developmental time points, n=5, **p<0.01, ***p<0.001 by One-way ANOVA with Fisher's LSD. **B-C.** Western blot quantification of MEX3B protein expression levels in mouse cortex at different developmental time points, n=5, *p<0.05, ***p<0.001, ****p<0.0001 by One-way ANOVA with Fisher's LSD. **D.** Comparison of normalized Mex3b protein and RNA levels in Mouse cortex at different developmental time points, n=5, *p<0.05, **p<0.01, ***p<0.001, ****p<0.0001 by Two-way ANOVA with Fisher's LSD. All the data is shown as Mean±SEM. Significance in A and C is represented with respect to E11.5 to rest of the developmental time points. **E-F.** GO analysis for Cellular component (E.) and Molecular function (F.) from Mex3b targets obtained from Microarray data from previous studies.

A.

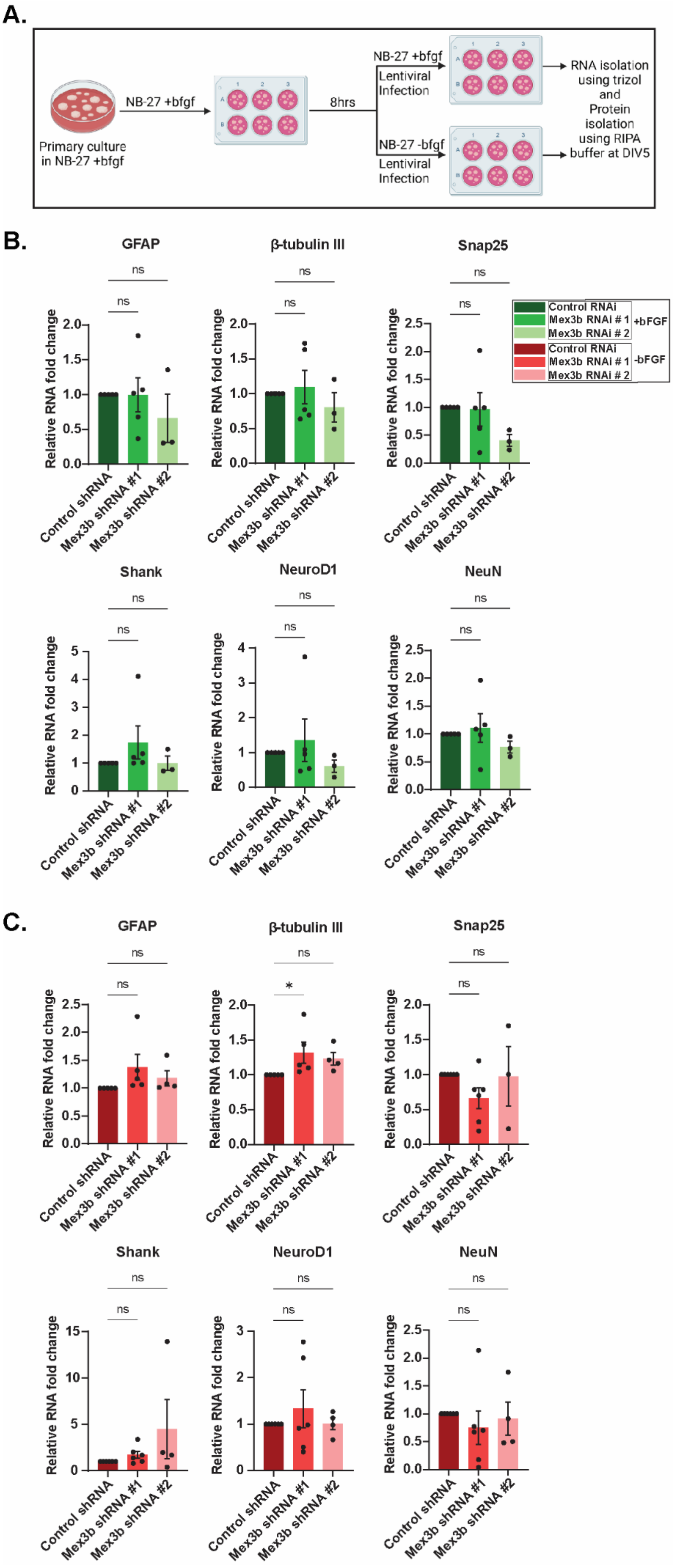
Primary culture in NB-27 +bfgf
NB-27 +bfgf
8hrs
NB-27 +bfgf
Lentiviral Infection
NB-27 -bfgf
Lentiviral Infection
RNA isolation using trizol and Protein isolation using RIPA buffer at DIV5
B.
GFAP
β-tubulin III
Snap25
Shank
NeuroD1
NeuN
ns
Relative RNA fold change
Control shRNA
Mex3b shRNA #1
Mex3b shRNA #2
Control RNAi
Mex3b RNAi # 1
Mex3b RNAi # 2
+bFGF
-bFGF
C.
*

**FigureS3: Ineffectiveness of MEX3B RNAi on transcript level of differential markers. A.** Schematics of lentiviral infection in NPCs. **B-C.** Differential markers transcriptional expression in plus-bFGF and minus-bFGF condition in NPCs. n=3-6, *p<0.05, ns= not significant by One-way ANOVA with Fisher's LSD. All the data is shown as Mean±SEM.

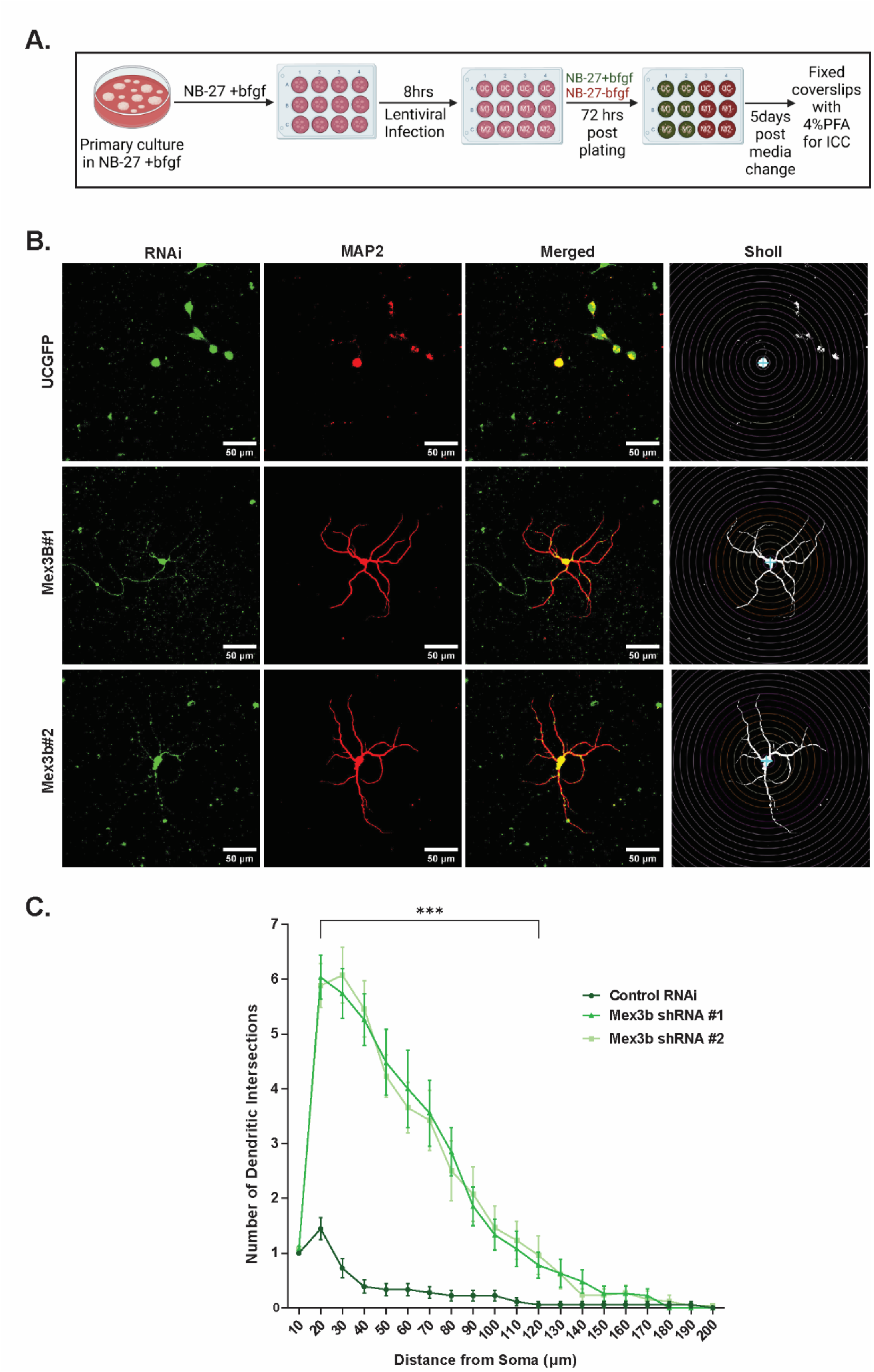


**FigureS4: MEX3B affects dendritic complexity. A.** Schematics of ICC on lentiviral infected NPCs. **B.** Photomicrographic representation for sholl analysis in NPCs. **C.** Quantification of number of dendritic intersections from soma. n=18-27 NPCs, ***p=

significant changes at distances from 10μm to 120μm by Two-way Anova with Multiple comparison at each point and Fisher's LSD test.

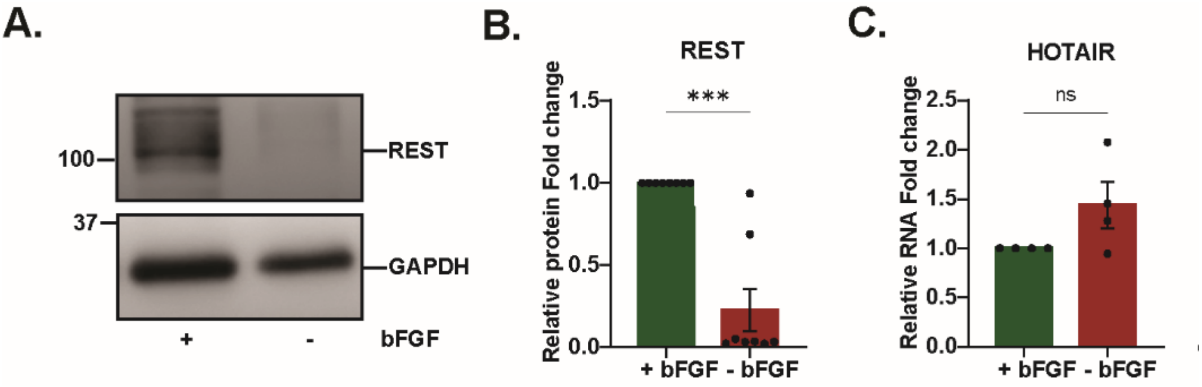


**FigureS5: REST and HOTAIR expression in NPCs. A-B.** Quantification of REST protein expression in differentiating vs proliferating NPCs. n=8, ***p<0.001. **C.** qRT-PCR based expression of HOTAIR lncRNA in differentiating vs proliferating NPCs. n=4, ns= not significant.
